# Systems-level network modeling of Small Cell Lung Cancer subtypes identifies master regulators and destabilizers

**DOI:** 10.1101/506402

**Authors:** David J Wooten, Sarah M Groves, Darren R Tyson, Qi Liu, Jing S Lim, Carlos F Lopez, Julien Sage, Vito Quaranta

## Abstract

Adopting a systems approach, we devise a general workflow to define actionable subtypes in human cancers. Applied to small cell lung cancer (SCLC), the workflow identifies four subtypes based on global gene expression patterns and ontologies. Three correspond to known subtypes, while the fourth is a previously undescribed neuroendocrine variant (NEv2). Tumor deconvolution with subtype gene signatures shows that all of the subtypes are detectable in varying proportions in human and mouse tumors. To understand how multiple stable subtypes can arise within a tumor, we infer a network of transcription factors and develop BooleaBayes, a minimally-constrained Boolean rule-fitting approach. In *silico* perturbations of the network identify master regulators and destabilizers of its attractors. Specific to NEv2, BooleaBayes predicts ELF3 and NR0B1 as master regulators of the subtype, and TCF3 as a master destabilizer. Since the four subtypes exhibit differential drug sensitivity, with NEv2 consistently least sensitive, these findings may lead to actionable therapeutic strategies that consider SCLC intratumoral heterogeneity. Our systems-level approach should generalize to other cancer types.

**Author summary:** Small-cell lung cancer (SCLC) is an extremely aggressive disease with poor prognosis. Despite significant advances in treatments of other cancer types, therapeutic strategies for SCLC have remained unchanged for decades. We hypothesize that distinct SCLC subtypes with differential drug sensitivities may be responsible for poor treatment outcomes.

To this end, we applied a computational pipeline to identify and characterize SCLC subtypes. We found four subtypes, including one (termed “NEv2”) that had not previously been reported. Across a broad panel of drugs, we show that NEv2 is more resistant than other SCLC subtypes, suggesting that this subtype may be partly responsible for poor treatment outcomes. Importantly, we validate the existence of NEv2 cells in both human and mouse tumors.

Reprogramming the identity of NEv2 cells into other subtypes may sensitize these cells to existing treatments. However, deciphering global mechanisms that regulate different subtypes is generally unfeasible. To circumvent this, we developed BooleaBayes, a modeling approach that only infers local regulatory mechanisms near stable cell subtypes. Using BooleaBayes, we found master regulators and master destabilizers for each subtype. These findings predict targets that may destabilize a particular subtype, including NEv2, and lead to successful therapy, by either knocking out master regulators or turning on master destabilizers.

## Introduction

A major barrier to effective cancer treatment is the occurrence of heterogeneous cell subpopulations that arise within a tumor via genetic or non-genetic mechanisms. Clonal evolution of these subpopulations via plasticity, drug-induced selection, or transdifferentiation allows tumors to evade treatment and relapse in a therapy-resistant manner. Characterizing cancer subpopulations, or subtypes, has led to breakthrough targeted treatments that significantly improve patient outcomes, as in the case of melanoma [1], breast [2], and lung cancer [3]. However, approaches to subtype identification suffer from several limitations, including: i) focus on biomarkers, which frequently possess insufficient resolving power; ii) lack of consideration for the system dynamics of the tumor as a whole; and iii) often phenomenological, rather than mechanistic, explanations for subtype sources.

To accelerate progress in cancer subtype identification, we set out to develop a general systems-level approach that considers underlying molecular mechanisms to generate multiple stable subtypes within a histological cancer type. We focused on gene regulatory networks (GRNs) comprised of key transcription factors (TFs) that could explain the rise, coexistence and possibly trans-differentiation of subtypes. To enumerate subtypes, identify key regulating TFs, and predict reprogramming strategies for these subtypes, we established the workflow shown in Fig 1. Briefly, we use consensus clustering and weighted gene co-expression network analysis on transcriptomics data to identify cancer subtypes distinguished by gene expression signatures, biological ontologies, and drug response. We validate the existence of the subtypes in both human and mouse tumors using CIBERSORT [4] and nearest neighbor analyses, and develop a GRN that can explain the existence of multiple stable subtypes within a tumor. We then introduce BooleaBayes, a Python-based algorithm to infer partially constrained regulatory interactions from steady state gene expression data. Applied to this GRN, BooleaBayes identifies and ranks master regulators and master destabilizers of each subtype. In a nutshell, starting from transcriptomics data, the workflow can predict reprogramming strategies to improve efficacy of treatment.

**Fig 1.**
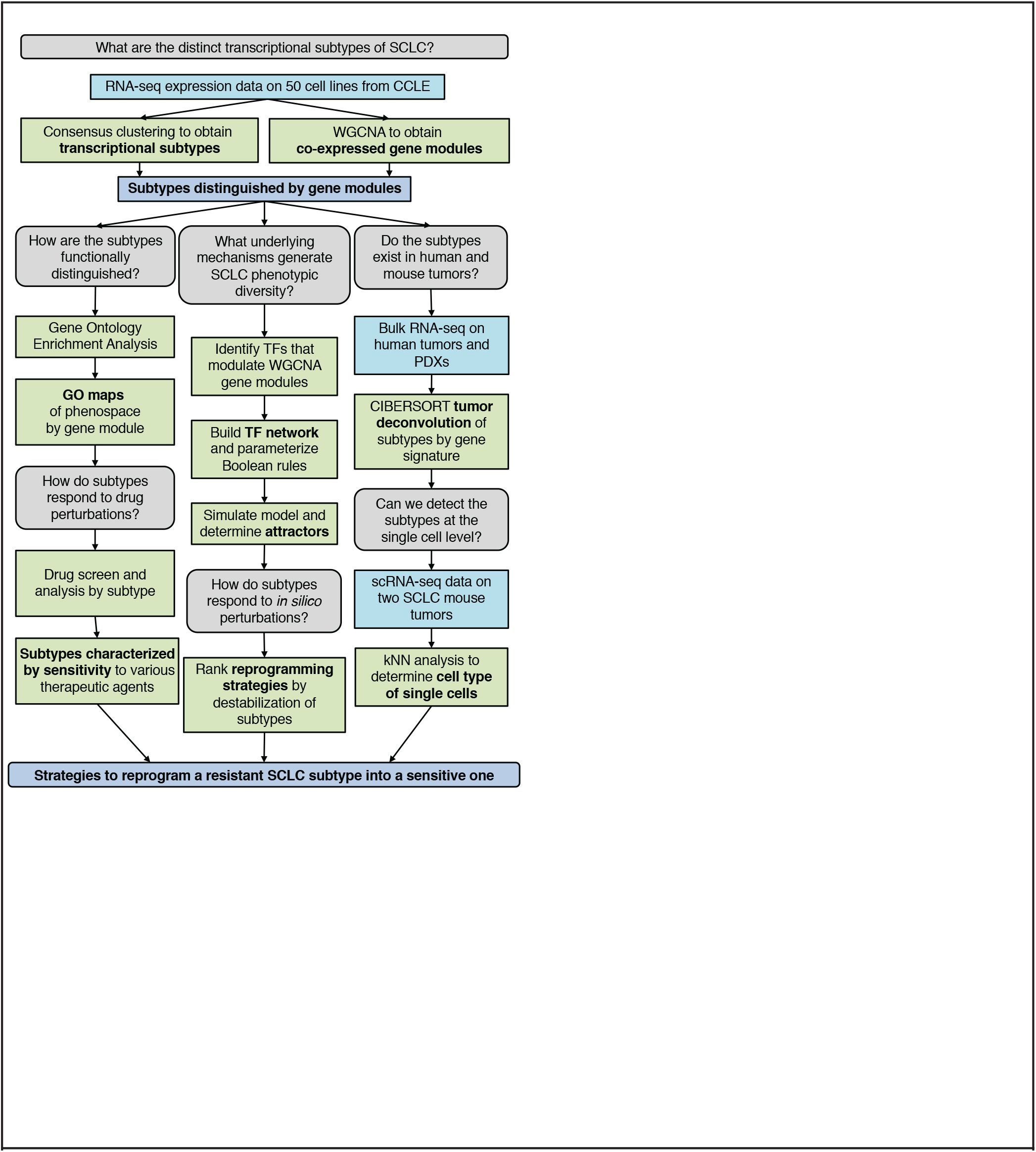
Workflow of our analysis. We use parallel analyses to identify strategies to reprogram resistant SCLC subpopulations into sensitive ones. These strategies can then be tested in *vitro* and *in vivo*.

We applied this workflow to Small Cell Lung Cancer (SCLC), in which genetic aberrations cannot fully distinguish subtypes [5], or point toward a targeted therapy. SCLC treatment has instead remained cytotoxic chemotherapy (a regimen of etoposide and a platinum-based agent such as cisplatin) and radiation for over half a century, despite the fact that virtually all patients relapse after therapy. This has caused SCLC to be designated as a recalcitrant cancer by the Recalcitrant Cancer Research Act of 2012, with five year survival rates less than 5%.

Recently, efforts to stratify patients have led to the recognition of phenotypic heterogeneity within and between SCLC tumors, raising hopes for more efficient subtype-based treatment strategies. As first described over 30 years ago, human SCLC cell lines can be categorized into two broad subtypes: a neuroendocrine (NE) stem-cell-like “classic” subtype and a distinct non-NE “variant” subtype [6–8]. In both human and mouse tumors, most cells appear to belong to the NE subtype, corresponding to a pulmonary neuroendocrine cell (PNEC) of origin [9], with high expression of neuroendocrine genes such as ASCL1. However, several groups have found evidence for non-NE variants within SCLC tumors [10–12], as well as an NE variant driven by MYC overexpression and NEUROD1 overexpression, instead of ASCL1 [13–15]. We previously described SCLC cell lines with hybrid expression of both NE and non-NE markers [16], and proposed they could serve as a resistant niche since drug perturbations shifted most cell lines towards hybrid phenotype(s). Taken together, these observations indicate the existence of a complex landscape of SCLC phenotypes that may form a tumor microenvironment robust to perturbations and treatment [10,17]. However, previous SCLC subtype reports were limited in their ability to systematically identify subtypes and understand plasticity across them. We hypothesized that our workflow, by taking into account the dynamics of underlying GRNs, could make systems-level predictions that more accurately reflect the occurrence and transdifferentiation of coexisting subtypes within SCLC tumors.

Starting from transcriptomics data from SCLC cell lines, our pipeline identifies four transcriptional subtypes, and a GRN that describes their dynamics. Three of these correspond to known ones, the fourth is a previously unreported NE variant (termed NEv2) with reduced sensitivity to drugs. Both CIBERSORT and single-cell validation reveal that in virtually every human and mouse tumor heterogeneity encompasses NEv2, and that all other previously reported subtypes are represented across tumors. BooleaBayes identifies both master regulators and master destabilizers for each subtype, opening the way for treatment strategies that may take SCLC subtypes into account. For instance, we hypothesize that by targeting these master TFs, the NEv2 phenotype may be destabilized leading to increased treatment sensitivity of SCLC tumors.

## Materials and methods

### Data

Human SCLC cell line data was taken from the Broad Institute’s CCLE RNA-seq expression data (version from February 14, 2018) at https://portals.broadinstitute.org/ccle/data. 81 human tumors were obtained from George et al. dataset, courtesy of R.K. Thomas [5]. The Myc-high mouse data set [15] was obtained from the NCBI GEO deposited at GSE89660. PDX/CDX mouse data [18] was obtained from the NCBI GEO deposited at GSE110853. Data was subsetted to only include SCLC cell lines (50). Features with consistently low read counts (< 10 in all samples) and non-protein-coding genes were removed. All expression data was then converted to TPM units and log1p normalized by dataset.

### Clustering and WGCNA

We applied Consensus Clustering to RNA-seq gene expression data from the 50 SCLC cell lines in the Cancer Cell Line Encyclopedia (CCLE) using the ConsensusClusterPlus R package [19]. Gene expression (TPM) was median-centered prior to clustering, and we clustered the cell lines using a k-means method with a Pearson distance metric for k ∈ {2,12} (S1 Fig). Other parameters were set as follows: reps = 1000, pItem = 0.8, pFeature = 0.8, seed = 1. Best k value was chosen heuristically based on the cumulative distributive function plot (S1 Fig, tracking plot, delta area plot, and average consensus scores (not shown).

To identify gene programs driving the distinction between the four SCLC phenotypic clusters, we performed weighted gene co-expression network analysis (WGCNA) on the same RNA-seq data. The softPower threshold was chosen as 12 to generate a signed adjacency matrix from gene expression. A topological overlap matrix (TOM) was created using this adjacency matrix as input. Hierarchical clustering on 1-TOM using method = ‘average,’ and the function cutTreeDynamic was used to find modules with parameters: deepSplit=2, pamRespectsDendro=TRUE, minClusterSize=100. These settings were chosen based on an analysis of module stability and robustness. We then computed an ANOVA comparing the four subtypes for each module. 11 out of 18 modules were able to statistically distinguish between the four clusters with an FDR-adjusted p-value < 0.05.

### Gene Ontology Enrichment Analysis

We ran a gene ontology (GO) enrichment analysis on each module that was significantly able to distinguish the phenotypes (11 total). The terms that were significantly enriched in at least one module were culminated into a general list of terms enriched in SCLC, which had 1763 terms. To visualize these terms, we computed a distance matrix between pairs of GO terms using GoSemSim [20], and used this matrix to project the terms into a low dimensional space using t-SNE. t-SNE is a popular method that computes a low-dimensional embedding of data points and seeks to preserve the high-dimensional distance between points in the low-dimensional space.

### Drug Sensitivity Analysis

Our drug sensitivity analysis used the freely available drug screen data from Polley, et al [21]. This screen included 103 Food and Drug Administration-approved oncology agents and 423 investigational agents on 63 human SCLC cell lines and 3 NSCLC lines. We subsetted the data to the 50 CCLE cell lines used for our previous analyses that had defined phenotypes according to Consensus Clustering (above). As described in [21], “the compounds were screened in triplicate at nine concentrations with a 96-hour exposure time using an ATP Lite endpoint.” Curve fitting, statistical analysis, and plotting was done by Thunor Web, a web application for managing, visualizing and analyzing high throughput screen (HTS) data developed by our lab at Vanderbilt University [22]. For fitting the dose response curve, as mentioned in the Thunor manual (https://docs.thunor.net/): “Thunor fits viability data to a three parameter log-logistic model. The three parameters are *E_max_, EC*_50_, and Hill coefficient. A constrained fit is used: *E_max_* is constrained to be between 0 and 1, and the Hill coefficient (slope) is constrained to be non-negative.” A measure of activity area for each drug-cell line combination was calculated as “the area above the observed response values, up to the no response value (1 on the y-axis). Between doses, a straight line extrapolation is used (on a log10 dose x-axis). Response values above 1 are truncated at 1. If there are multiple response values for a particular concentration (replicates), the mean average is used.” This takes into account both the efficacy and potency of a drug when added to each cell line. By segregating the cell lines by subtype, we were able to evaluate the relationship between drug response and subtype. Further information is available at https://docs.thunor.net/.

## CIBERSORT

CIBERSORT is a computational inference tool developed by Newman et al. at Stanford University [4]. We utilized the interactive user interface of CIBERSORT Jar Version 1.06 at https://cibersort.stanford.edu/runcibersort.php. Gene signatures were automatically determined by the software from a provided sample file with a matching phenotype class file. For this sample file and class file, the RNA-seq data from 50 human SCLC cell lines were inputted with their consensus clustering class labels. For each run, 500 permutations were performed. Relative and absolute modes were run together, with quantile normalization disabled for RNA-seq data, kappa = 999, q-value cut-off = 0.3, and 50-150 barcode genes considered when building the signature matrix.

### Single cell RNA sequencing of TKO SCLC tumors

The p53, Rb and p130 triple-knockout (TKO) SCLC mouse model with the Rosa26membrane-Tomato/membrane-GFP (Rosa26mT/mG) reporter allele has been described (Denny and Yang et al., 2016). Tumors were induced in 8-weeks old TKO; Rosa26mT/mG mice by intratracheal administration of 4×107 PFU of Adeno-CMV-Cre (Baylor College of Medicine, Houston, TX). 7 months after tumor induction, single tumors (one tumor each from two mice) were dissected from the lungs and digested to obtain single cells for FACS as previously described [10,23]. DAPI-negative live cells were sorted using a 100 *μ*m nozzle on a BD FACSAria II, spundown and resuspended in PBS with 10% bovine growth serum (Fisher Scientific) at a concentration of 1000 cells/*μ*1. Single cell capture and library generation was performed using the Chromium Single Cell Controller (10x Genomics) and sequencing was performed using the NextSeq High-output kit (Illumina).

### Single cell analysis

Cells with ≤ 500 detected genes per cell or with ≤ 10% of transcripts corresponding to mitochondria-encoded genes were removed. Low abundance genes that were detected in less than 10 cells were excluded. Each cell was normalized to a total of 10,000 UMI counts, and log2-transformed after the addition of 1. Top 1000 highly variable genes were selected and clusters of cells were identified by the shared nearest neighbor modularity optimization based on the top 10 PCs using the highly variable genes and visualized by t-SNE in R package Seurat [24]. The k-nearest neighbors (kNN) with k=10 of human cell lines was detected for each mouse cell to predict subtypes of the individual cell based on signature genes of each subtype. If at least 80% nearest human cell line neighbors for a mouse cell belong to one subtype, the mouse cell was assigned to that subtype. Otherwise, the subtype was undetermined (not assigned).

### Genomic analysis

Mutational Analysis was performed by MutSigCV V1.2 from the Broad Institute [25]. First, a dataset of merged mutation calls (including coding region, germ-line filtered) from the Broad Cancer Dependency Map [26] was subsetted to only include SCLC cell lines. Background mutation rates were estimated for each gene-category combination based on the observed silent mutations for the gene and non-coding mutations in the surrounding regions. Using a model based on these background mutation rates, significance levels of mutation were determined by comparing the observed mutations in a gene to the expected counts based on the model. MutSigCV was run on the GenePattern server using this mutation table, the territory file for the reference human exome provided for the coverage table file, the default covariate table file (gene.covariates.txt), and the sample dictionary (mutation_type_dictionary_file.txt). Only genes with an FDR-corrected q-value < 0.25 were considered significant, which are shown in S2 Fig.

### BooleaBayes inference of logical relationships in the TF network

A Boolean function of *N* input variables is a function *F* : {0,1}^N^ ↦ {0,1}. The domain of *F* is a finite set with 2^N^ elements, and therefore *F* is completely specified by a 2^*N*^ dimensional vector in the space {0,1}^2^ in which each component of the vector corresponds to the output of *F* for one possible input. In general, knowledge of the steady states of *F* is unlikely to be sufficient to fully constrain all 2^*N*^ components of the vector describing *F*. BooleaBayes is a practical approach that constrains *F* in the neighborhood of stable fixed points based on steady-state gene expression data. In practice, we let each component of the vector be a continuous real-value *v_i_* ∈ [0,1] reflecting our confidence in the output of *F*, based on available constraints. Components of *F* that are near 0.5 will indicate uncertainty about whether the output should be 0 or 1, given the available constraining data.

Given *M* observations (in our case, each observations is a measurement of gene expression of the *N* regulator TFs and the target TF in *M* = 50 cell lines), we want to compute this vector 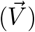 describing a probabilistic Boolean function *F* of *N* variables (see (5) in Fig 7B). First, we organize the input-output relationship as a binary tree with *N* layers leading to the 2^*N*^ leaves (see (2) in Fig 7B), each of which corresponds to a component of vector 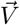. For instance, given two regulators A and B (*N* = 2), the leaves of the binary tree correspond to the probabilities that 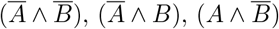, and 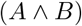. Collectively, the observations define an *M* × *N* matrix 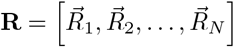 quantifying the input regulator variables (columns) for each observation (rows), as well as an *M* dimensional vector 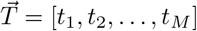 quantifying the output variable (see (1) in Fig 7B). A Gaussian mixed model is then used to transform the columns of **R** (regulator variables) and the vector 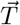 into probabilities **R**′ and 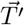 of the variables being OFF or ON in each observation (row).

Let 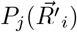 be a function that quantifies the probability that the input variables of the *i*^th^ observation belong to the *j*^th^ leaf of the binary tree. For instance using the example above, the second leaf of the binary tree is 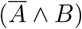. Therefore, *P*_*j*=2_(*A, B*) = (1 – *A*) · *B*. Note that by this definition, 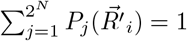. Using this, we define an *M* × 2^*N*^ weight matrix **W** = *w_i,j_* as

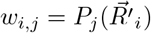

that describes how much the *i*^th^ observation constrains the *j*^th^ component of 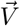 (see the grayscale heatmap in Fig 7B). Additionally, to avoid overfitting under-determined leaves (columns in the grayscale heatmap with no dark cells, meaning that none of the observations constrain that leaf), we define the uncertainty 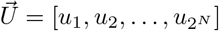 of each leaf

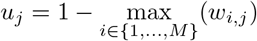

From these, we then define the vector 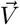 describing function *F* as

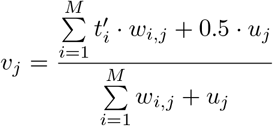

Thus, each component of 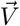 is the average of the output target variable 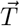 weighted by **W**, with an additional uncertainty term 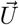 to avoid overfitting. For leaves *j* of the binary tree that are poorly constrained by any of the observables, *v_j_* ≈ 0.5, indicating maximal uncertainty in the output of *F* at those leaves. Uncertainty of a leaf *j* also arises when observations *i* with large weight *w_i_ j* have inconsistent values for 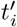, such as if 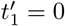 and 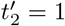.

## Results

### Consensus Clustering uncovers new SCLC variant phenotype

Recently, the occurrence of variant SCLC subtypes has been reported [13,15,16]. Given the translational value of defining subtypes, a more global approach to comprehensively define SCLC subytpes would be desirable. To this end, we devised the workflow described in Fig. 1. First, we applied Consensus Clustering [27] to RNA-seq gene expression data from the 50 SCLC cell lines in the Cancer Cell Line Encyclopedia (CCLE) [28]. We clustered the cell lines using a k-means method with a Pearson distance metric for k ∈ {2, 20} (Fig 2A). Both k=2 and k=4 gave well defined clusters. Since recent literature suggests that more than two subtypes are necessary to adequately describe SCLC phenotypic heterogeneity, we selected k=4 for further analyses.

**Fig 2.**
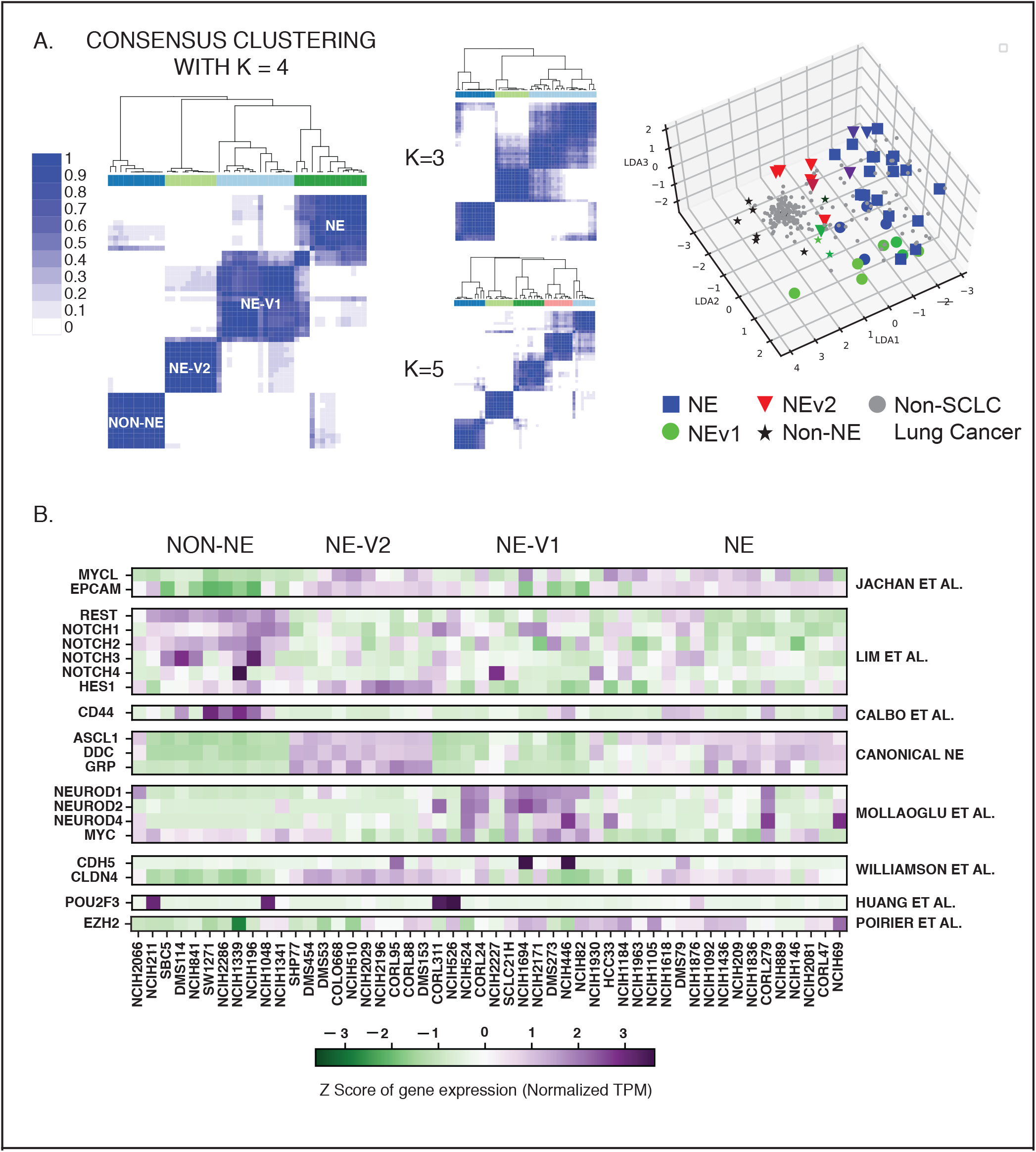
Consensus clustering and WGCNA of 50 SCLC cell lines reveals four subtypes differentiated by gene modules. **A.** Consensus clustering with k = 4 gives most consistent clusters. K = 3 and K=5 add complexity without a corresponding increase in accuracy. LDA plot shows separation of 4 clusters, with non-SCLC cell lines falling near non-NE cell lines. **B.** Current biomarkers in the field of SCLC are able to distinguish between three of the subtypes; The fourth subtype, NEv2, is not separable from NE using markers from SCLC literature.

To align the 4-cluster classification (Fig 2B) with existing literature, we considered distribution of well-studied biomarkers of SCLC heterogeneity across the clusters. Three of the four consensus clusters could be readily matched to subtypes previously identified with 2 to 5 biomarkers: the canonical NE subtype, an NE variant subtype (referred to here as NEv1), and a non-NE variant subtype [10,14,15,29]. However, the fourth cluster (referred to here as NEv2) could not be easily resolved using only these few markers. For example, NEv2 may be considered a tumor propagating cell (TPC) by biomarkers in Jachan et al [23], yet expression of *HES1* may suggest grouping into a non-NE subtype by Lim et al [10]. This discrepancy drove us to consider broader patterns of gene expression, rather than a limited number of biomarkers, to characterize each subtype.

### SCLC phenotypes are differentially enriched in diverse biological processes, including drug catabolism and immuno-modulation

To capture global gene expression patterns, we applied Weighted Gene Co-expression Network Analysis (WGCNA) [30] to RNA-seq data from CCLE for multiple SCLC cell lines (See methods). This analysis revealed 17 groups, or modules, of co-expressed genes. Module eigengenes could be used to describe trends of gene expression levels. 11 of these 17 groups of co-expressed genes could statistically distinguish between the four consensus clusters (Fig 3A, Table S1, Kruskal-Wallis, FDR-adjusted p < 0.05). To specify the biological processes enriched in each of these 11 gene modules, we performed gene ontology (GO) enrichment analysis using the Consensus Path Database [31], which resulted in a combined total of 1,763 statistically enriched biological processes (Table S2, Fig 3B). In particular, the yellow, salmon, and pink modules are enriched for neuroendocrine differentiation and neurotransmitter secretion and are upregulated in the canonical NE and NEv1 phenotypes (Fig 3C). In contrast, the blue, black, and purple modules, enriched for cell adhesion and migration processes, are upregulated in the non-NE variant phenotype, in agreement with the observed adherent culture characteristics of these cell lines.

**Fig 3.**
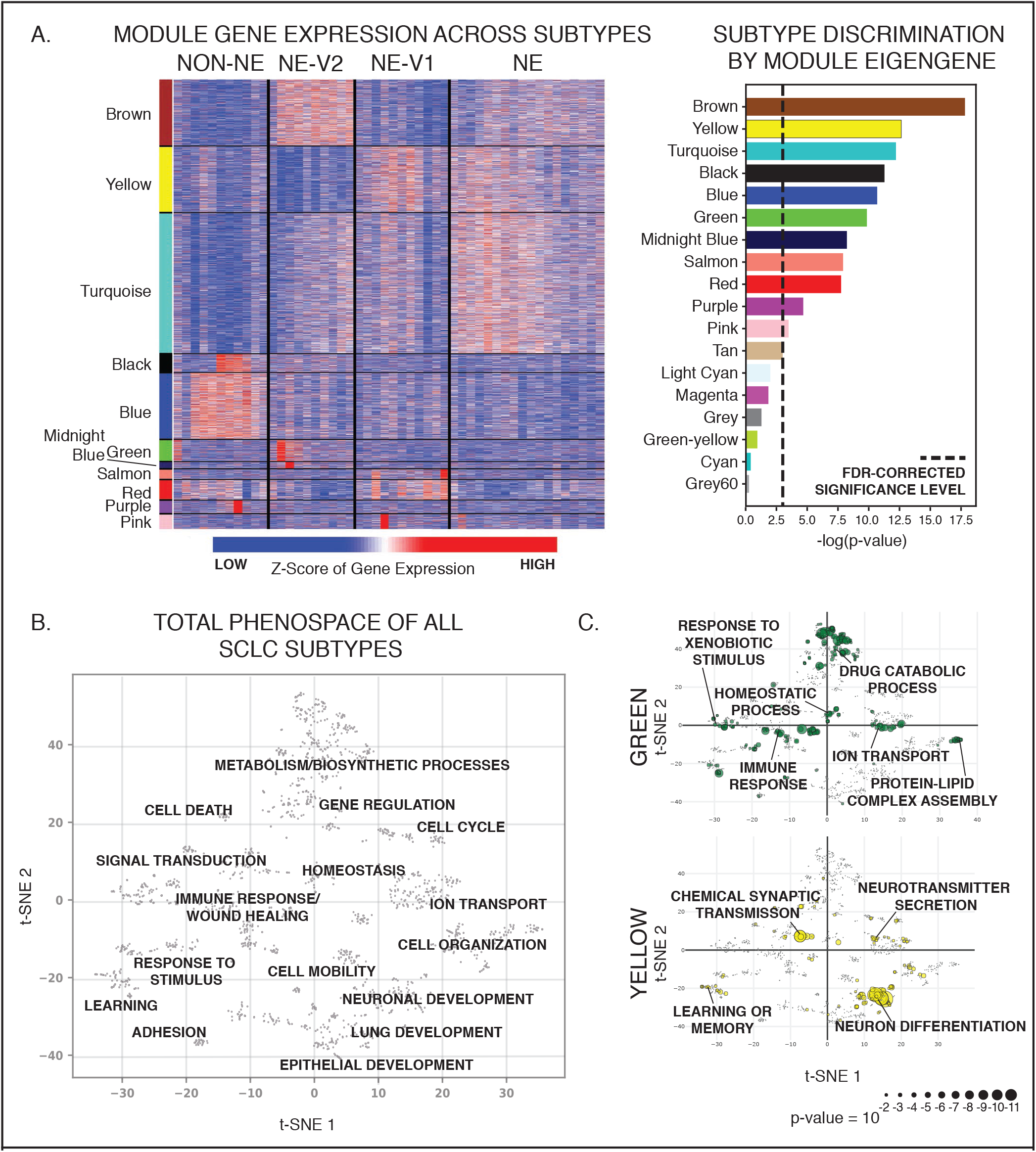
SCLC subtypes can be distinguished by gene expression patterns. **A.** Transcriptional patterns that distinguish the four subtypes are captured in WGCNA analysis. Gene modules by color show patterns of expression that are consistent across the subtypes. Only modules that significantly distinguish between the subtypes are shown (ANOVA, FDR-corrected p-value < 0.05). **B.** SCLC heterogeneity biological process phenospace. A dissimilarity score between pairs of SCLC-enriched GO terms was calculated using GoSemSim, and used to create a t-SNE projection grouping similar biological processes together. Each blue dot is a GO term, with selected terms highlighted. Several distinct clusters of related processes can be seen. **C.** Module-specific phenospace. A breakout of where some of the 11 statistically significant WGCNA modules fall in the GO space from A. Of particular interest, the green module, which is highly upregulated in the NEv2 phenotype, is enriched in metabolic ontologies, including drug catabolism and metabolism and xenobotic metabolism. The yellow module is enriched in canonical neuronal features.

Genes within the brown, midnight blue, and green modules are upregulated in the NEv2 phenotype. The brown module is enriched for canonical phenotypic features of SCLC, particularly cellular secretion and epithelial differentiation, and accordingly is also upregulated in the canonical NE subtype. The midnight blue module, enriched in nervous system processes and lipid metabolism, is highly expressed in the NEv2 cell lines. The green module is enriched for immune/inflammatory response, wound healing, homeostasis, drug/xenobiotic metabolism, and cellular response to environmental signals (Fig 3C). Enrichment of these GO terms suggest that NEv2 cells may more easily adapt to external perturbations such as therapeutic agents, and potentially show higher drug resistance.

To visualize these enriched GO terms in an organized way (Fig 3B), we used the GOSemSim package [20] in R to compute a pairwise dissimilarity score, or distance, between all enriched GO terms (FDR-adjusted p < 0.05 in at least one of the 11 significant modules). We then projected all significant GO terms into a 2D space by t-distributed stochastic neighbor embedding (t-SNE) [32]. In this t-SNE projected phenospace, GO terms that describe semantically similar biological processes are placed close to one another and grouped into a general biological process. This map allows exploration of biological processes enriched in individual gene modules or subtypes, and it shows that SCLC heterogeneity spans biological processes that can largely be grouped as 1) related to neuronal, endocrine, or epithelial differentiation; 2) metabolism and catabolism; 3) cell-cell adhesion and mobility; and 4) response to environmental stimuli, including immune and inflammatory responses. In summary, the phenospace constructed from global gene expression patterns captures the unique characteristics of each SCLC subtype.

### Drug resistance is a feature of the NEv2 subtype

As mentioned previously, the enriched GO terms for drug catabolism and xenobiotic metabolism in the green module suggest that the NEv2 phenotype may have a higher ability to metabolize drugs and therefore exhibit decreased sensitivity. To test this possibility, we reanalyzed drug responses of SCLC cell lines to a panel of 103 FDA-approved oncology agents and 423 investigational agents in the context of our four subtype classification [21]. We used the Activity Area (AA) metric as a measure of the resultant dose-response curves. The drugs were analyzed individually and clustered by common mechanism of action and target type, and the cell lines were grouped by the four subtypes (Table S3). Across all evaluated drugs, the NEv2 subtype exhibited the most resistance (54% cell lines were resistant). In contrast, both NE and NEv1 exhibited less resistance (20% cells were resistant), with non-NE exhibiting the least resistance (6% of cell lines were resistant) (Fig 4A). Taken together, these results confirm that based on the prediction from the gene-regulation based classification, the subtypes exhibit different levels of resistance and that high resistance is a feature of the NEv2 subtype (Fig 32C). In particular, mTOR inhibitors are a class of compounds to which NEv2 was significantly more resistant (Fig 4). PI3K pathway mutations have previously been implicated as oncogenic targets for SCLC, as about a third of patients show genetic alterations in this pathway [33]. Among the four subtypes, NEv2 is also the least sensitive to AURKA, B, and C inhibitors (AURKA shown); TOPO2 inhibitors; and HSP90 inhibitors (Fig 4). These results have implications for interpreting expected or observed treatment response with respect to tumor heterogeneity in individual patients.

**Fig 4.**
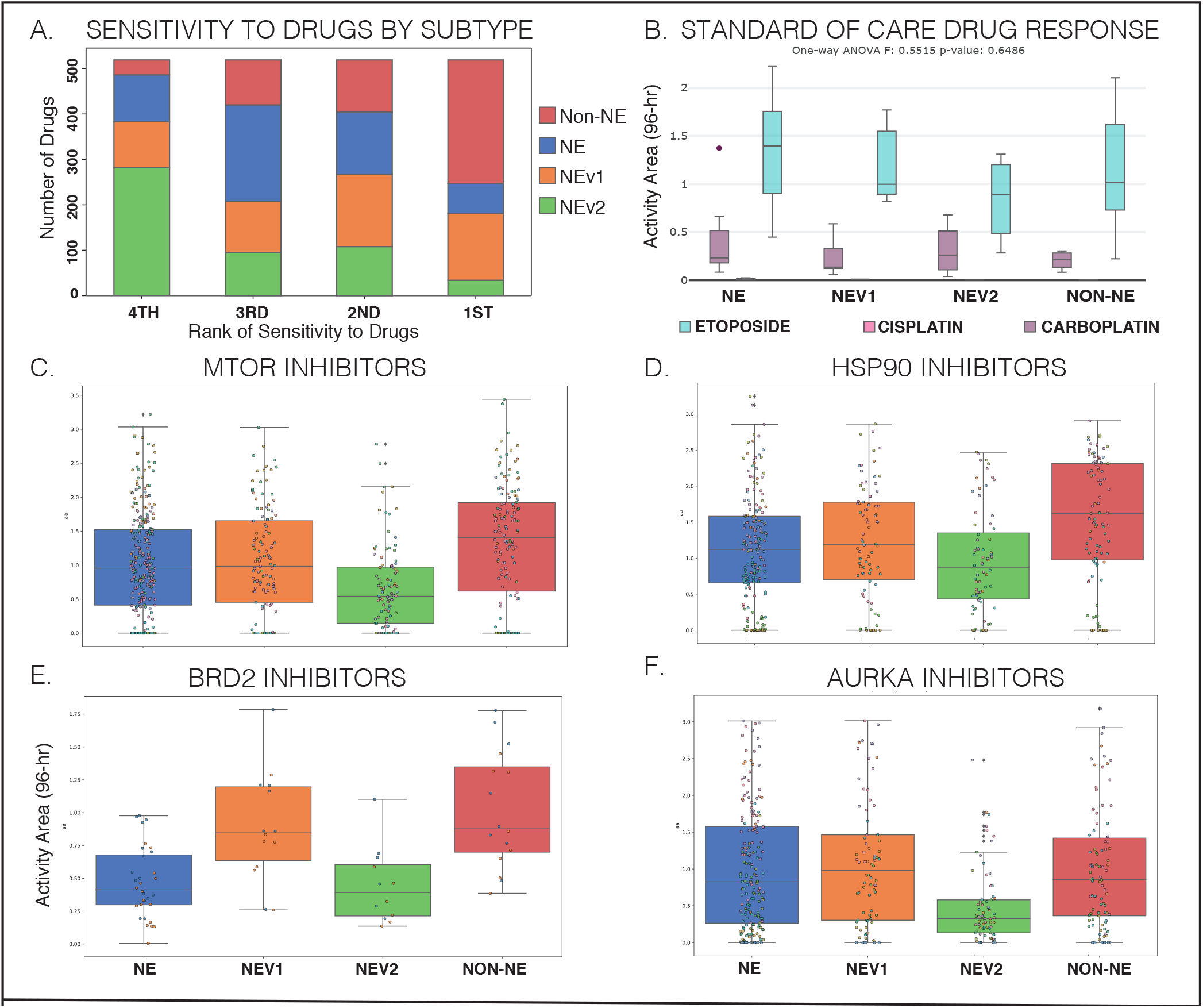
Differential response of SCLC subtypes to a wide variety of oncology drugs and investigational agents. **A.** Ranked sensitivity of subtypes across 526 compounds. NEv2 is least sensitive for over half of the drugs tested. **B.** No significant differences can be seen in response to etoposide and platinum-based agents cisplatin and carboplatin, the standard of care for SCLC. **C-F.** Significantly differential response by ANOVA, p < 0.05, shown in drugs that target **C.** mTOR, **D.** HSP90, **E.** BRD2, and **F.** AURKA. NEv2 is significantly more resistant to all of these drugs.

### Neuroendocrine variants are represented in mouse and human SCLC tumors

Next, we investigated whether the four subtypes we detected in human SCLC cell lines are also present in tumors. We used CIBERSORT [4] to generate gene signatures for each of the 4 subtypes. These gene signatures could then deconvolve RNA-seq measurements on 81 SCLC tumors from George et al [5] to specify relative prevalence of each subtype within a single tumor. CIBERSORT predicted that a majority of tumors were comprised of all four subtype signatures, in varying proportions across tumor samples (Fig 5A). We then analyzed the patient/cell-derived xenograft models (PDXs/CDXs) developed by Drapkin et al [18], and the tumors also showed vast differences across samples (Fig 5B). Some of these samples were taken across multiple time points from the same patient, thus enabling us to test both tumor composition and dynamic changes in tumor subpopulations. Three samples taken from patient MGH1514, before and after treatment, indicated a change in tumor composition in favor of the NE phenotype. In contrast, patient MGH1518 showed a reduction of NEv1 and increase in NEv2 after treatment. Overall, the high variance in proportions of each subtype suggest a high degree of intertumoral, as well as intratumoral, dynamic heterogeneity and plasticity.

**Fig 5.**
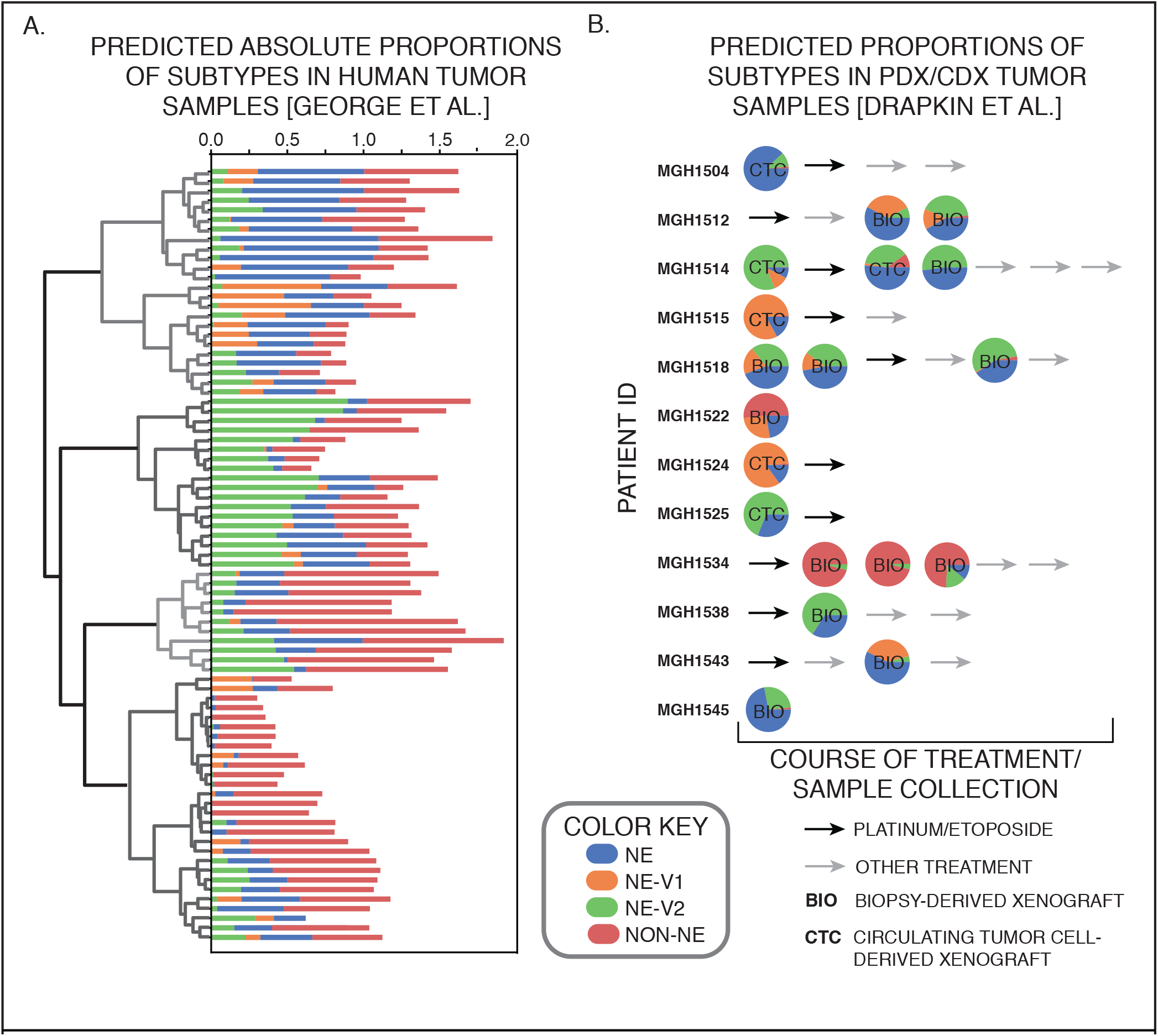
Computational evidence for existence of subtypes in human tumors. **A.** Absolute percentage of each subtype in 81 human tumors as determined by CIBERSORT. The 81 tumors can then be sorted by hierarchical clustering, which finds four main groups of subtype patterns across tumors. **B.** Similar analysis in mouse PDX/CDX tumors from Drapkin et al. [18]. As shown, the tumors vary greatly in composition, and hierarchical clustering of the patterns result in four clusters (clustering not shown).

We also investigated phenotypic patterns in mouse tumors from two different sources to determine whether human SCLC subtype signatures are conserved across species [15] (Supplemental Methods). The first mouse model is a triple knockout *(Rb1, Tp53*, and *P130*, conditionally deleted in lung cells via a Cre-Lox system, TKO), and these tumors were primarily composed of the NE and NEv2 subtypes (Fig 6Ai). Of note is the lower percentage of non-NE cells found in each tumor in Fig 6Ai; we suspect this is due to a filtering step before sequencing (Supplemental Methods), as the non-NE subtype signature is more similar to tumor-associated immune cells in an unfiltered tumor population. The second mouse model was generated with *Myc* overexpression (double knockout of *Rb1* and *Tp53*, and overexpression of MYC) (Fig 6Aii) as reported previously [15]. Using the subtype gene-signatures developed in the previous sections, the Myc-high tumors showed a clear increase in the percentage of NEv1 detected compared to the triple knockout tumors in Fig 6Ai, corroborating the correlation between NEv1 and a previously described Myc-high mouse tumor subtype.

**Fig 6.**
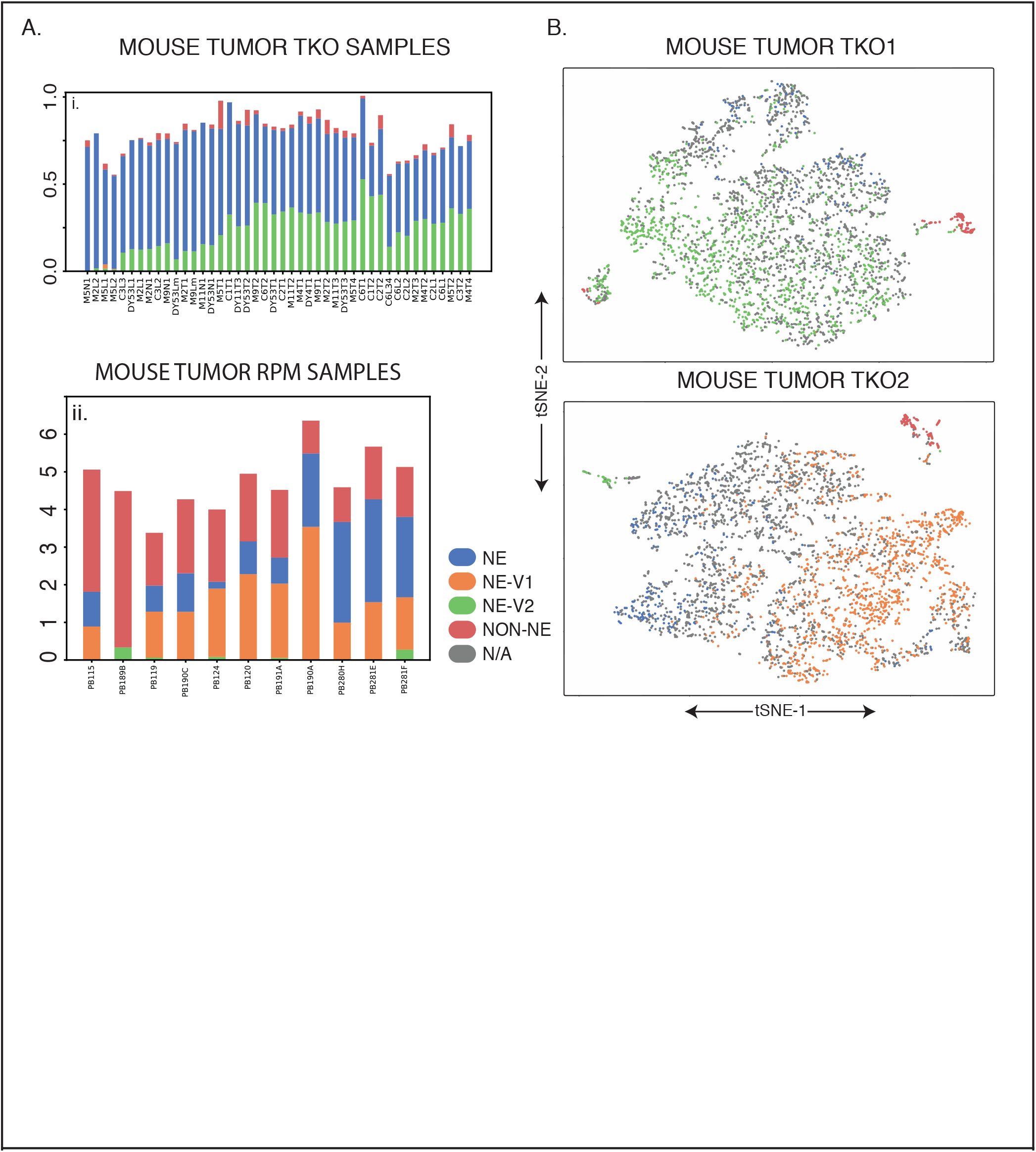
Similar analysis in mouse tumors. **A.** Ai. TKO *(Rb1, Tp53, P130* floxed) mouse tumors showing a high proportion of NE and NEv2 subtypes. Aii. As described in [15], these mouse tumors were generated by crossing Rb1 fl/fl Trp53 fl/fl (RP) animals to knockin Lox-Stop-Lox (LSL)-MycT58A-IRES-Luciferase mice. These Rb1 fl/fl Trp53 fl/fl Myc LSL/LSL (RPM) mice have overexpressed Myc, and have been shown to be driven towards a variant phenotype, which is corroborated in this CIBERSORT analysis. It is clear that RPM mice contain greater portions of NEv1 compared to the tumors in Ci., which seems to correspond to the Aurora-Kinase-inhibitor-sensitive, *Myc*-high phenotype published by Mollaoglu et al. **B.** t-SNE plots of single cell RNA-seq from two TKO mouse tumors. The k-nearest neighbors (kNN) with k=10 was computed for each mouse cell to predict subtypes of individual cell using signature genes of each subtype. If at least 8 of the 10 nearest human cell line neighbors for a mouse cell were of one subtype, the cell was assigned that subtype. Large amounts of intratumoral and intertumoral heterogeneity are evident.

Lastly, we analyzed two primary TKO mouse tumors by single cell RNA-seq (scRNA-seq). For each mouse single cell transcriptome, we computed the k=10 nearest *human cell line* neighbors (kNN with k=10), and assigned each mouse cell to a subtype based on its neighbors (Supplemental methods). As shown in Fig 6B, a large portion of the cells from each tumor correspond to one of the four human subtypes. A small non-NE population can be seen in both tumors, and about a third of the assigned cells correspond to the NE subtype (Fig 5B). Tumor A has a large proportion of the NEv2 subtype, corresponding to the tumors in 5Ai. In contrast, tumor B has a large NEv1 subpopulation, similar to the tumors in 5Aii. Taken together, these results indicate that subtypes in SCLC tumors are conserved across species, and can be categorized either by CIBERSORT analysis of bulk transcriptomics data, or by kNN analysis of scRNA-seq data.

### Genetic mutations alone cannot account for four SCLC phenotypes

The evidence above for intratumoral and intertumoral heterogeneity led us to investigate how the subtypes arise and coexist in both human and mouse SCLC tumors. To determine whether mutations could be responsible for defining the four SCLC subtypes, we analyzed genomic data in the Broad Cancer Dependency Map [26]. We subsetted these data to the 50 SCLC cell lines with matching CCLE RNA-seq data, and using MutSigCV [25], we found 29 genes (S2 Fig) mutated more often than expected by chance (using a significance cutoff of q-value ≤ 0.5 to be as inclusive as possible). However, none of these genes were able to separate the four subtypes by mutational status alone (S2 Fig), suggesting alternative sources of heterogeneity.

### Transcription factor network defines SCLC phenotypic heterogeneity and reveals master regulators

To investigate these alternative sources of heterogeneity, we hypothesized that different SCLC subtypes emerge from the dynamics of an underlying TF network. We previously identified a TF network that explained NE and non-NE SCLC subtype heterogeneity [16]. That analysis suggested the existence of additional SCLC subtypes but did not specify corresponding attractors [16]. Here, we performed an expanded TF network analysis to find stable attractors for all four SCLC subtypes. As an initial step, we identified putative master TF regulators within each of the 11 WGCNA modules (Fig 3B) based on differential expression. Regulatory interactions between these TFs were extracted from public databases, including ChEA, TRANSFAC, JASPAR, and ENCODE, based on evidence of TF-DNA binding sites in the promoter region of a target TF, as well as several sources from the literature. This updated network largely overlaps with, but contains several refinements compared to our previous report [16], as detailed in Fig 7A.

**Fig 7.**
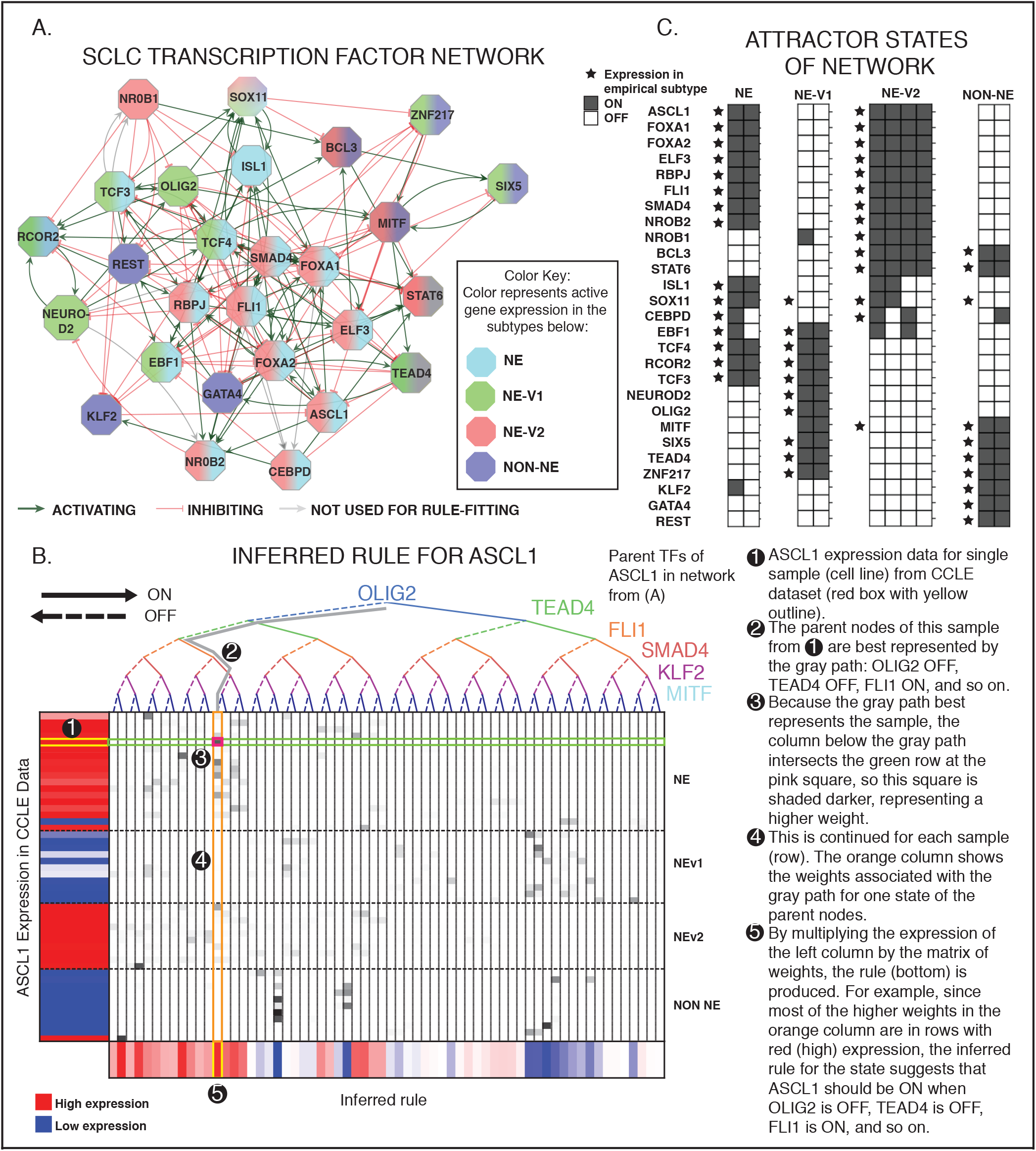
TF network simulations reproduce subtypes as attractors. **A.** Regulatory network of differentially expressed TFs from each of the 11 co-expressed gene modules in Fig 2B. Colors indicate which phenotype each TF is upregulated in. Red edges indicate inhibition (on average), and green activation (on average). **B.** Probabilistic Boolean rule fits for ASCL1. The target gene is a function of all the genes along the binary tree at the top, while expression of the target is shown on the left. Each row represents one cell line, each column represents one possible input state, and the bottom shows the inferred function F for every possible input state. Color ranges from 0=blue (highly confident the TF is off), to 0.5 = white, to 1=red (highly confident the TF is on). Rows are organized by subtype (top to bottom: NE, NEv1, NEv2, non-NE). **C.** Attractors found with asynchronous updates of Boolean network. 10 attractors were found, and each correlates highly with one of the four subtypes (represented by stars). Specifics of the probabilistic simulation are described in Results.

Following the procedure we previously used [16], we simulated the network using threshold and inhibitory dominant update rules. These update rules are commonly used as coarse-grained approximations of systems for which detailed biological regulatory functions are unknown. However, these approximations were not sufficient to stabilize attractors corresponding to either the NEv1 or NEv2 phenotypes (data not shown), suggesting that the regulatory rules governing stability of these phenotypes are more complex.

To understand this, we developed BooleaBayes, a method to infer logical relationships in gene regulatory networks (Fig 7B) by enhancing confidence in Boolean rules via a Bayes-like adjustment approach (Supplemental methods). With BooleaBayes, we were able to describe the stabilization of all four SCLC phenotypes from steady-state gene expression data (Fig 7C). An advantage of this method is that it makes predictions intrinsic to those parts of the network in which we are most confident (Supplemental Methods).

BooleaBayes rules (probabilistic logical relationships between connected nodes) were derived for each node of the SCLC TF network in Fig 7A. As an example, Fig 7B shows the rule fitting for one node, ASCL1. Rules for all other nodes are given in Supplemental Fig 4. We simulated the dynamics of the Boolean network using a general-asynchronous update scheme [34]. This formed a state transition graph (STG), in which each state is defined by a vector of TF ON/OFF expression values.

For the initial states in the simulation, we discretized the average TF expression for each of the four SCLC subtypes. We exhaustively searched the neighborhood of each of these starting states out to a distance of 6 TF changes in the STG. Within these neighborhoods, we found 10 states for which all 27 TFs had at least a 50% chance of remaining unchanged. Transitions into these states are therefore more likely, and escapes less likely. Thus, these 10 states represent semi-stable states of the network dynamics (Fig 7C), that we refer to as pseudo-attractors.

These 10 pseudo-attractor states each correlated with, and could be assigned to, one of the 4 SCLC subtypes (stars in Fig.4C); this indicates that the updated network structure and BooleaBayes rules are sufficient to capture stability of the four SCLC phenotypes. Having identified network dynamics that closely match experimental observations, we are now in a position to perform *in silico* (de)stabilizing perturbations and predict the resulting trajectory through the STG for each subtype. We do so in the next section.

### In *silico* SCLC network perturbations identify master regulators and master destabilizers of SCLC phenotypes

To quantify the baseline stability of the steady states in Fig 7C, we performed random walks (algorithm described in Supplemental Methods) starting from each of the 10 pseudo-attractors. We counted how many steps were required to reach a state outside of a 4-TF neighborhood around the starting state (Fig 8). We chose a 4-TF neighborhood for a given attractor because a randomly chosen state has only a 2.5% chance of falling within this neighborhood, suggesting states in this neighborhood are significantly similar to the attractor state (p = 0.025). Each TF in the network was either activated (held constant at TF=1) or silenced (TF=0) in each of the stable states, and 1000 random walks were executed for each condition (Fig 8A). The percent increase or decrease of the stability relative to the unperturbed reference was calculated, resulting in a score quantifying (de)stabilization of the starting state by each TF perturbation (Fig 8B, Fig S3). For example, either activation of GATA4 (TF = 1) or silencing FOXA1 (TF=0) are predicted to destabilize both the NE and NEv2 subtypes (Fig 8C).

**Fig 8.**
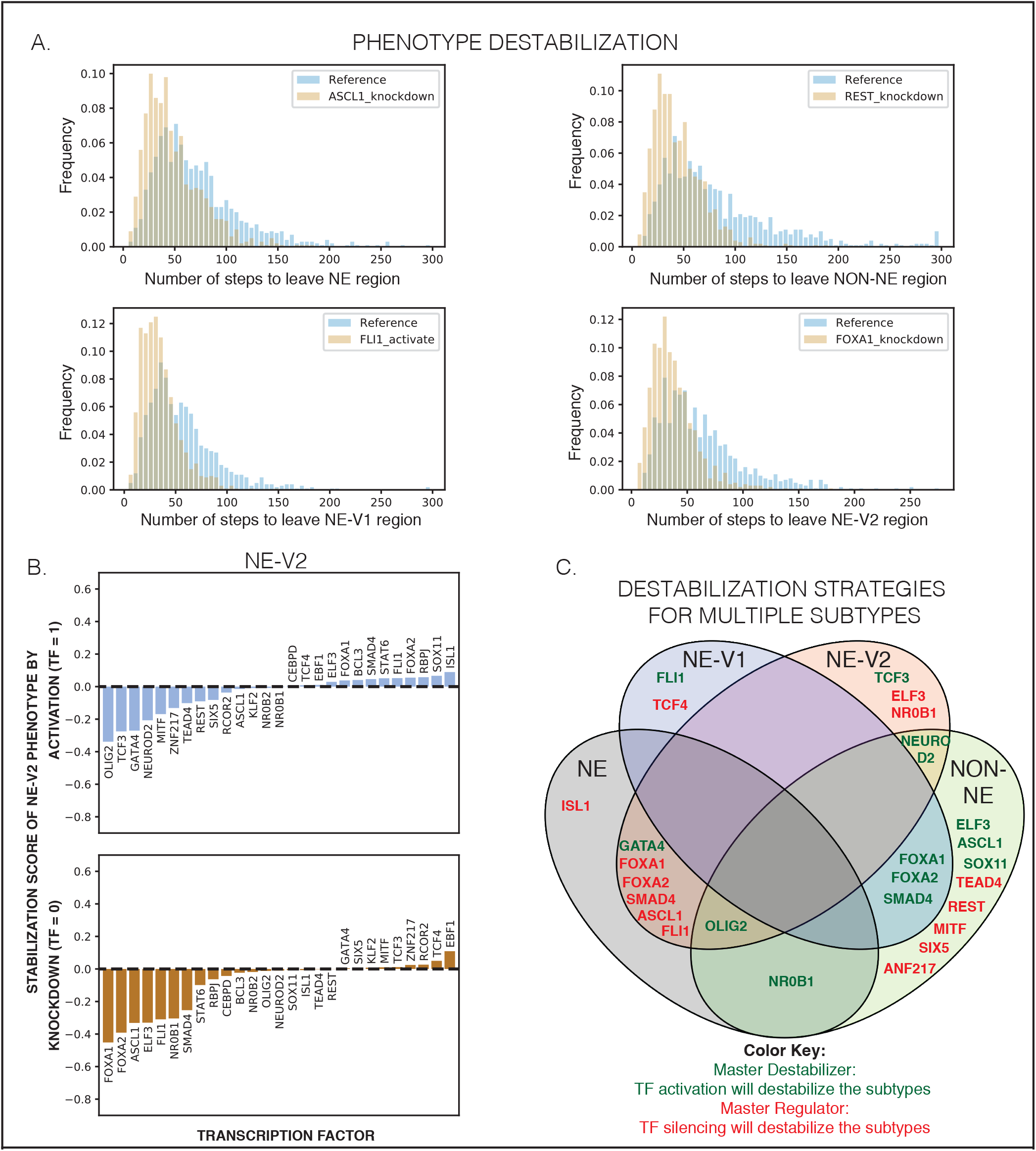
Destabilization of subtypes by perturbation to network. **A.**Random walks starting from the attractors in Fig 4C will eventually leave the start state due to uncertainty in the Boolean rules. Reference histogram shows how many random steps are required for a distance greater than 4 TF changes away from the start state under the network’s natural dynamics. The knockdowns and activations shown here hold expression of the perturbed gene OFF or ON. The perturbation destabilizes the start state, such that the random walk leaves the neighborhood sooner, shown for NE, NEv1, NEv2, and non-NE starting states. **B.** Stabilization of SCLC phenotype NEv2 by TF activation and knockdown. The percent change of stability measures the percent change in the average number of steps needed to leave the neighborhood of the stable states. Negative indicates destabilizing, while positive indicates increasing stability. Results are shown for 1000 iterations starting from NEv2. Similar plots for the other subtypes can be found in Fig S3. **C.** A Venn diagram demonstrating overlap of destabilization strategies. A single activation (green text) or knockdown (red text) can sometimes destabilize multiple phenotypes. TF perturbations that have a stabilization score of less than −0.2 were considered destabilizing.

TFs that, when silenced, cause destabilization greater than 20% (score ≥ −0.2) of a specific subtype were considered master regulators of that subtype. They include *REST* (non-NE) (in agreement with [10], *TEAD4* (non-NE), *ISL1* (NE), and *TCF4* (NEv1). TEAD4 is downstream mediator of YAP1 action, which has been previously identified as a possible phenotypic modulator in a subset of SCLC cell lines [35]; our analyses suggest that expression of TEAD4 may be able to stabilize this phenotype. Simulations of the network also identified the novel NEv2 master regulators, *ELF3* and *NR0B1*.

Our network simulations further identified TFs that can be considered master “destabilizers”, i.e., *activation* of these TFs destabilizes a specific phenotype by at least 20%. For instance, activation of ELF3 is predicted to destabilize non-NE, while activation of NR0B1 would destabilize both non-NE and NE subtypes. Simulations identified a single master destabilizer for NEv2, the TF *TCF3* (Fig 8C). Taken together, our pipeline, which includes subtype identification, drug response analysis, and network simulations, suggests possible therapeutic perturbations that could shift the phenotypic landscape of SCLC into a more sensitive state for treatment.

## Discussion

We report a systems approach to understanding SCLC heterogeneity that integrates transcriptional, mutational, and drug-response data. Our findings culminate in discrimination and mechanistic insight into the four SCLC subtypes shown in Table 1: NE, non-NE, NEv1, and NEv2. Within the context of the broader literature on SCLC heterogeneity, we showed that NE, non-NE, and NEv1 correspond to several subtypes that have been previously reported based on a few markers. Significantly, we find that one (NEv2) has not been described previously, and which is nearly indistinguishable from NE based on currently used markers of SCLC heterogeneity.

**Table 1.** Comprehensive framework for SCLC heterogeneity. Using our workflow, we have characterized 4 SCLC subtypes by gene expression, drug response, and master regulators and destabilizers.

| Subtype | GO Terms | Gene Modules | Drug Response | Regulators | Destabilizers |
| --- | --- | --- | --- | --- | --- |
| NE | Neuron differentiation, Synaptic transmission, Cell-cycle regulation, Epithelial cell differentiation, Sensory perception | Turquoise, Brown, Yellow |  | ISL1, ASCL1, FLI1, SMAD4, FOXA1, FOXA2 | GATA4, OLIG2, NR0B1 |
| NEv1 | Neuron differentiation, neurotransmitter secretion, Synapse organization, Cardiac muscle cell contraction, Mechanoreceptor differentiation, Sensory perception | Yellow, Salmon, Pink, Red | Sensitive to AKIs | TCF4 | FLI1, FOXA1, FOXA2, SMAD4 |
| NEv2 | Immune response, Drug catabolism, Ion transport, Homeostasis, Epithelial cell differentiation, Glycosylation | Brown, Green, Midnight blue | Least sensitive | ELF3, NR0B1, FOXA1, FOXA2, SMAD4, ASCL1, FLI1 | TCF3, GATA4, NEUROD2, OLIG2 |
| NON-NE | Immune Response, Stress Response, Defense response, Macrophage activation, Myeloid cell differentiation and activation, neutrophil-mediated immunity, vesicle-mediated transport, Angiogenesis, Viral processes | Black, Blue, Red | Most sensitive | TEAD4, REST, MITF, SIX5, ZNF217 | ELF3, ASCL1, SOX11, FOXA1, FOXA2, SMAD4, NR0B1, OLIG2 |

Tumor deconvolution by CIBERSORT and analysis of scRNA-seq data indicate that this subtype is present in a large proportion of human and mouse SCLC tumors. Moreover, a drug screen across a broad range of compounds indicated that this subtype is more resistant than the others, especially in response to *AURK* and *mTOR* inhibitors. Direct experimental verification in mouse and human tumors of the NEv2 subtype’s drug resistant properties will be an important next step. For example, the PDXs from patient MGH1518 before and after drug show an increase in the NEv2 signature (Fig 5B), which may be responsible for acquired drug-resistance in this patient. However, this study was under-powered for our analyses, and more experimental data will be necessary to strengthen this conclusion.

A significant advance of our work is the introduction of BooleaBayes, which we developed to infer mechanistic insights into regulation of the heterogeneous SCLC subtypes. By considering the distinct subtype clusters as attractors of a gene regulatory network, BooleaBayes infers partially-constrained mechanistic models. A key benefit of this method is that it does not overfit data: predictions are based only on parts of the network for which available data can constrain the dynamics, while states that lack constraining data diffuse randomly. With this method we were able to recapitulate known master regulators of SCLC heterogeneity, as well as identify novel ones such as *ISL1* (NE) and *TEADĄ* (non-NE). Additionally, we predict *ELF3* and *NR0B1* to be master regulators of the NEv2 phenotype. Furthermore, we introduce the label of “master destabilizers” to describe TFs whose activation will destabilize a phenotype. Our method gives a systematic way to rank perturbations that may destabilize a resistant phenotype. In ongoing work, we are validating these predictions experimentally. We propose that with BooleaBayes, our approach for identifying master TFs could be applicable to other systems, including other cancer types or transcriptionally-regulated diseases.

While many of the previously reported subtypes of SCLC fit into our framework, a few are noticeably absent, and will require further study. The vasculogenic subtype of SCLC described by Williamson et al. [36] did not emerge from our analysis. We speculate that this may be due to the rarity and/or instability of this CTC-derived phenotype among the available SCLC cell lines. Denny and Yang et al. have previously reported that *Nfib* amplification promotes metastasis [37]; however, our clusters do not correlate with location of the tumor sample from which each cell line was derived (*e.g*., primary vs metastatic, data not shown). Poirier et al., using a similar clustering approach to ours, identified highly methylated SCLC subtypes (M1 and M2) [29], and the correspondence of these subtypes with the ones described here is intriguing and remains to be defined. Finally, Huang et al. recently reported an SCLC subtype defined by expression of *POU2F3* [12]. In our data, POU2F3 was highly expressed in only four cell lines, and was placed POU2F3 into a small (328 genes, green-yellow) module, and therefore represented only a small signal in our data. Overall, future studies with additional cell line and/or mouse data may be used to further investigate these different subtypes, underscoring that the delineation of four subtypes here does not preclude the existence of others.

## Conclusion

An advantage of our analyses is that each subtype is defined by distinct co-expressed gene programs, rather than by expression of one or few markers, which has been customary in the field but has limited ability to discriminate between phenotypes (Fig 2B). In addition, these modules participate in unique biological processes (e.g., as identified by GO), such that the systems-level approach presented here may provide a comprehensive framework to understand the regulation and functional consequences of SCLC heterogeneity in a tumor. This understanding can be actionable since SCLC subtypes show differential drug sensitivity; for example, our analyses in this paper support the hypothesis that NEv2 may be a drug-resistant phenotype of SCLC. We propose that identification of drugs targeting the NEv2 subtype, or perturbagens that reprogram it toward less recalcitrant states, may lead to improved treatment outcomes for SCLC patients.

## Supporting information

Pink GO Map

Salmon GO Map

Purple GO Map

Midnight Blue GO Map

Red GO Map

Black GO Map

Turquoise GO Map

Yellow GO Map

Blue GO Map

Brown GO Map

Green GO Map

## Supporting information

**S1 Fig.**
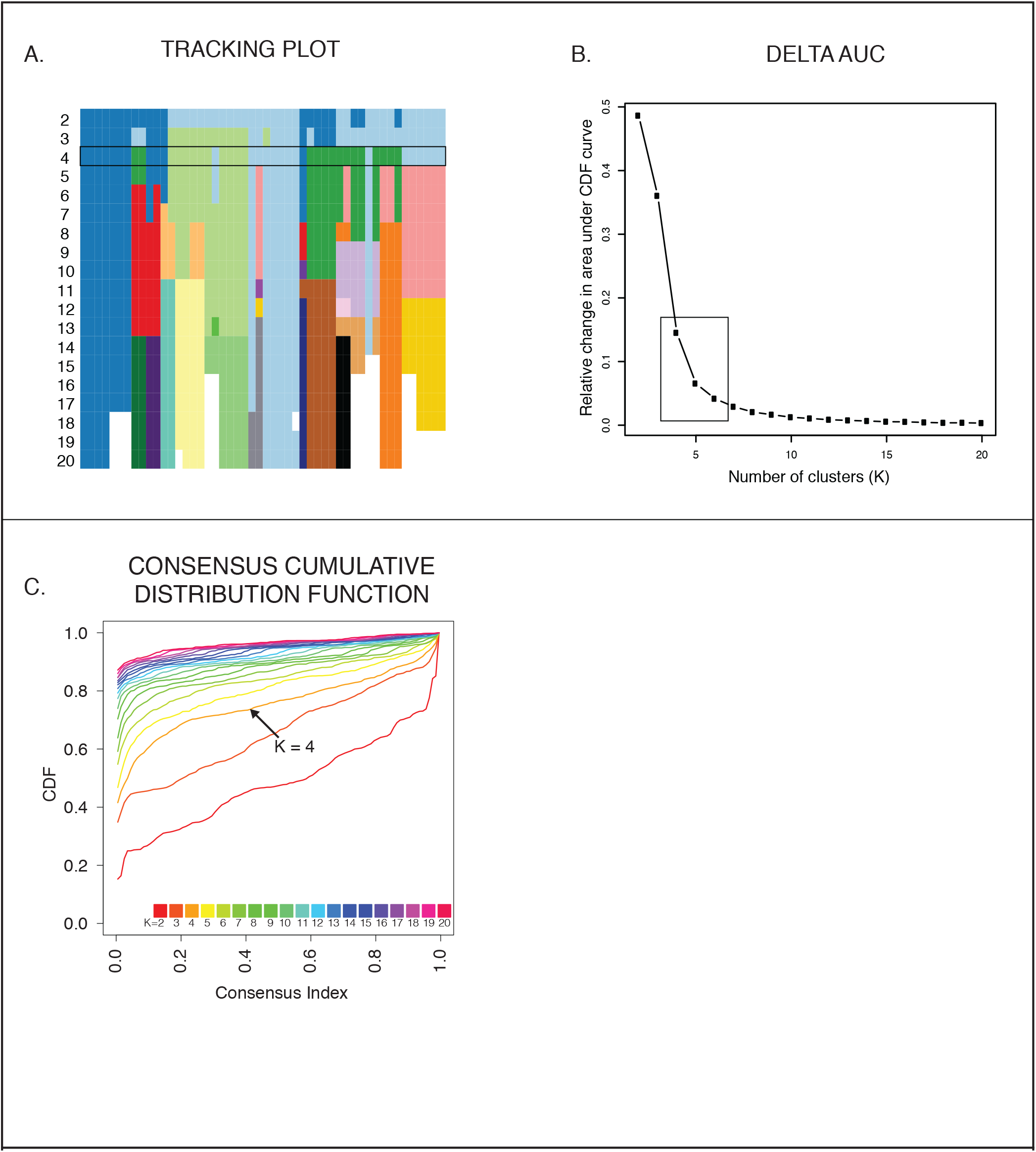
Tracking plot and delta area plot for consensus clustering with different values of k. **A.** Tracking plot shows slight inconsistency for cell lines with k=3. One of these is assigned to the “light green” cluster in the k=3 clustering scheme, whereas when k=4, it returns to the “light blue” cluster. The others are in the “dark blue” cluster when k=2 and “light blue” cluster when k=3. **B.** The delta area plot shows the relative change in the area under the CDF curve (Fig 1A). The largest changes in area occur between k=2 and k=4, at which point the relative increase in area becomes noticeably smaller (from an increase of 0.5 and 0.4 to 0.15). This suggests that k=4,5, or 6 are the best clustering that maximizes detail (more, smaller clusters present a more detailed picture than a few large clusters) and minimizes noise (by minimizing average pairwise consensus values and maximizing extreme pairwise consensus values. Average cluster consensus scores (CCS) across clusters show that k=4 may be the best choice because it has the highest average (k = 4 average CCS: 0.848, k=5 average CCS: 0.814, k=6 average CCS: 0.762). **C.** Consensus Cumulative Distribution Function. This CDF show that k=4 has more black cells and white cells than gray, suggesting the consensus clusters are more robust.

**S2 Fig.**
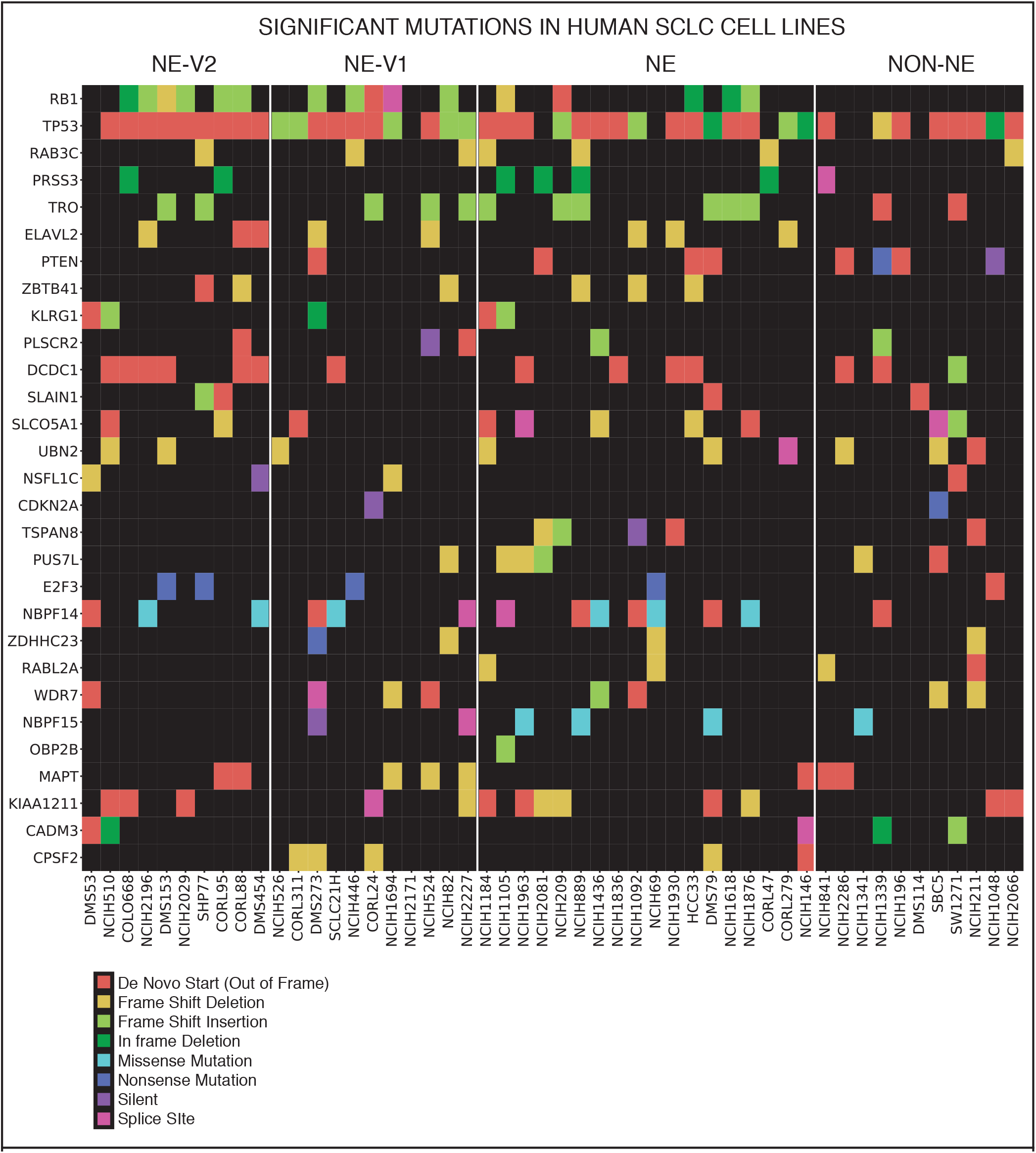
Significant mutations across subtypes. Significantly mutated genes across 50 SCLC cell lines, as determined by MutSigCV, ordered by significance. As expected, significant mutations were found in both the *Rb1* and *Tp53* genes. Inspection by eye shows that no significant mutations can distinguish completely between two or more phenotypes. This suggests an alternate source of heterogeneity, such as transcriptional regulation. Significance cut-off: q (p-value corrected for multiple comparisons) ≤ 0.25. q ≤ 0.5 shown.

**S3 Fig.**
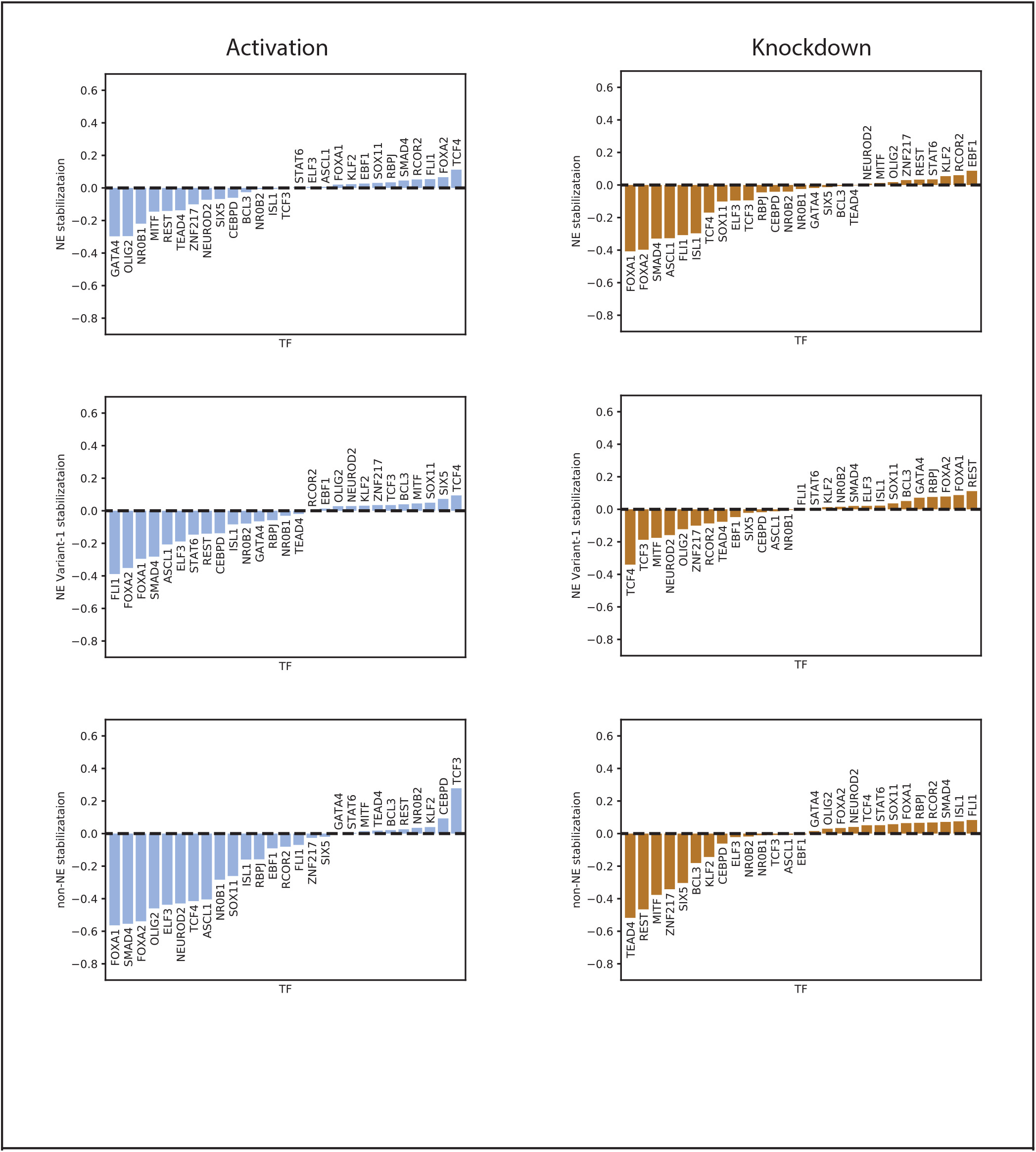
Stabilization of SCLC phenotypes by TF knockdown and activation. The percent change of stability measures the percent change in the average number of steps needed to leave the neighborhood of the stable states. Negative indicates destabilizing, while positive indicates increasing stability. Results are shown for 1000 iterations starting from **A.** NE, **B.** NEv1, and **C.** non-NE.

**S4 Fig. GO maps for each gene module.**

**S5 Fig. Rules for all transcription factors in network.**

**S1 Files. Interactive GO Maps.**

**S1 Table. Module Eigengenes.**

**S2 Table. Enriched Gene Ontology Terms in Subtype Modules.**

**S3 Table. Drug Response of Subtypes Grouped by Drug Target.**

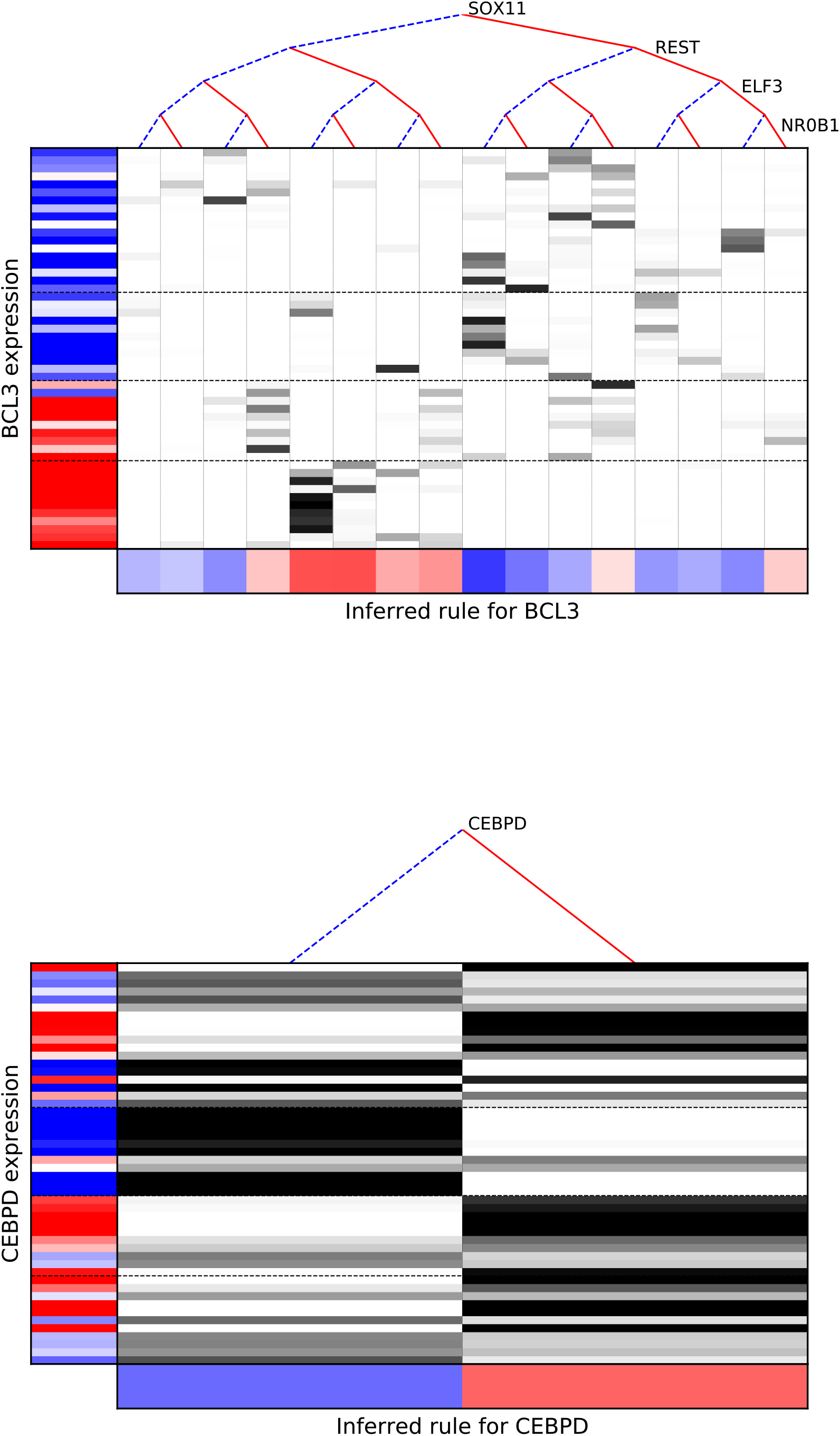

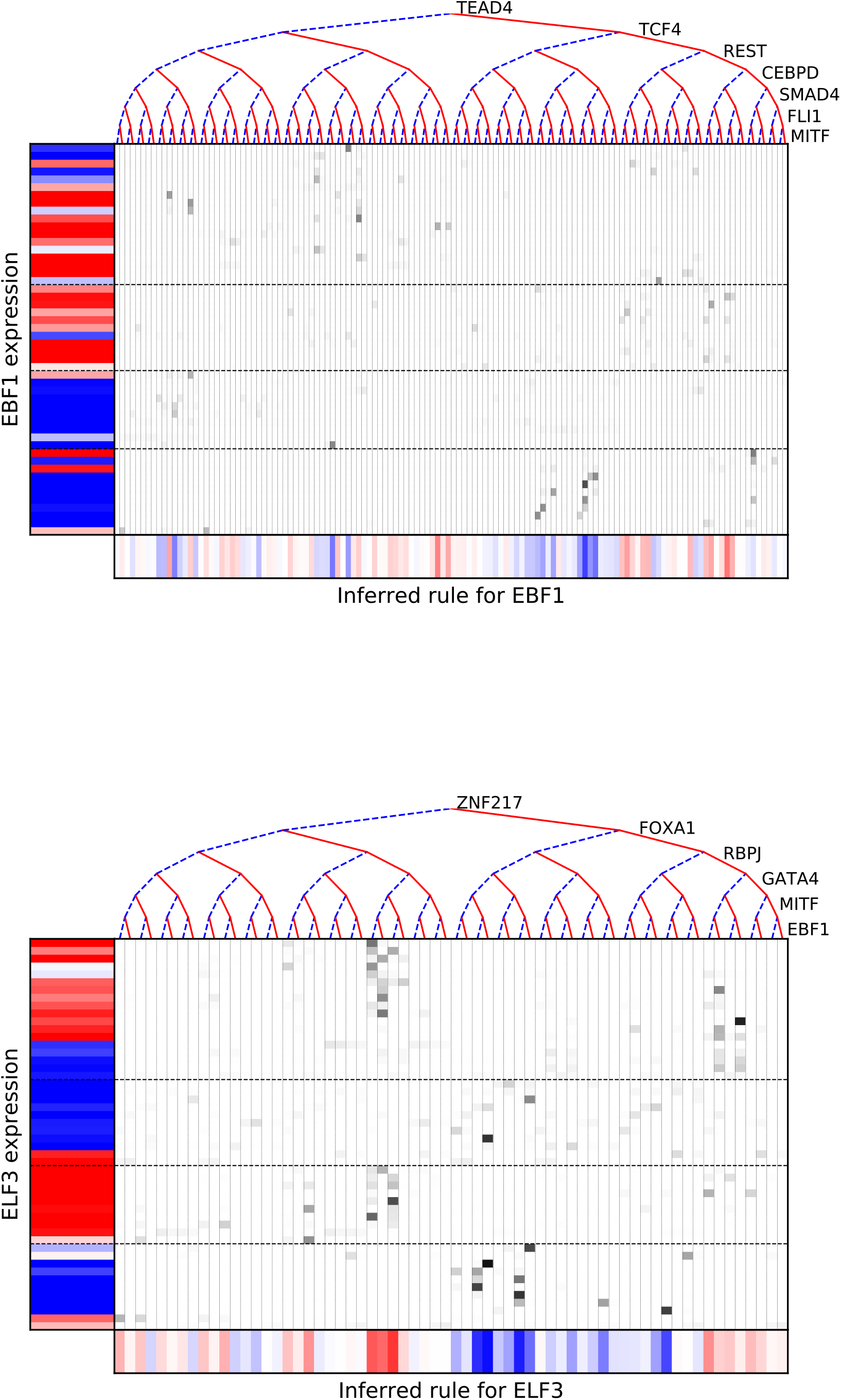

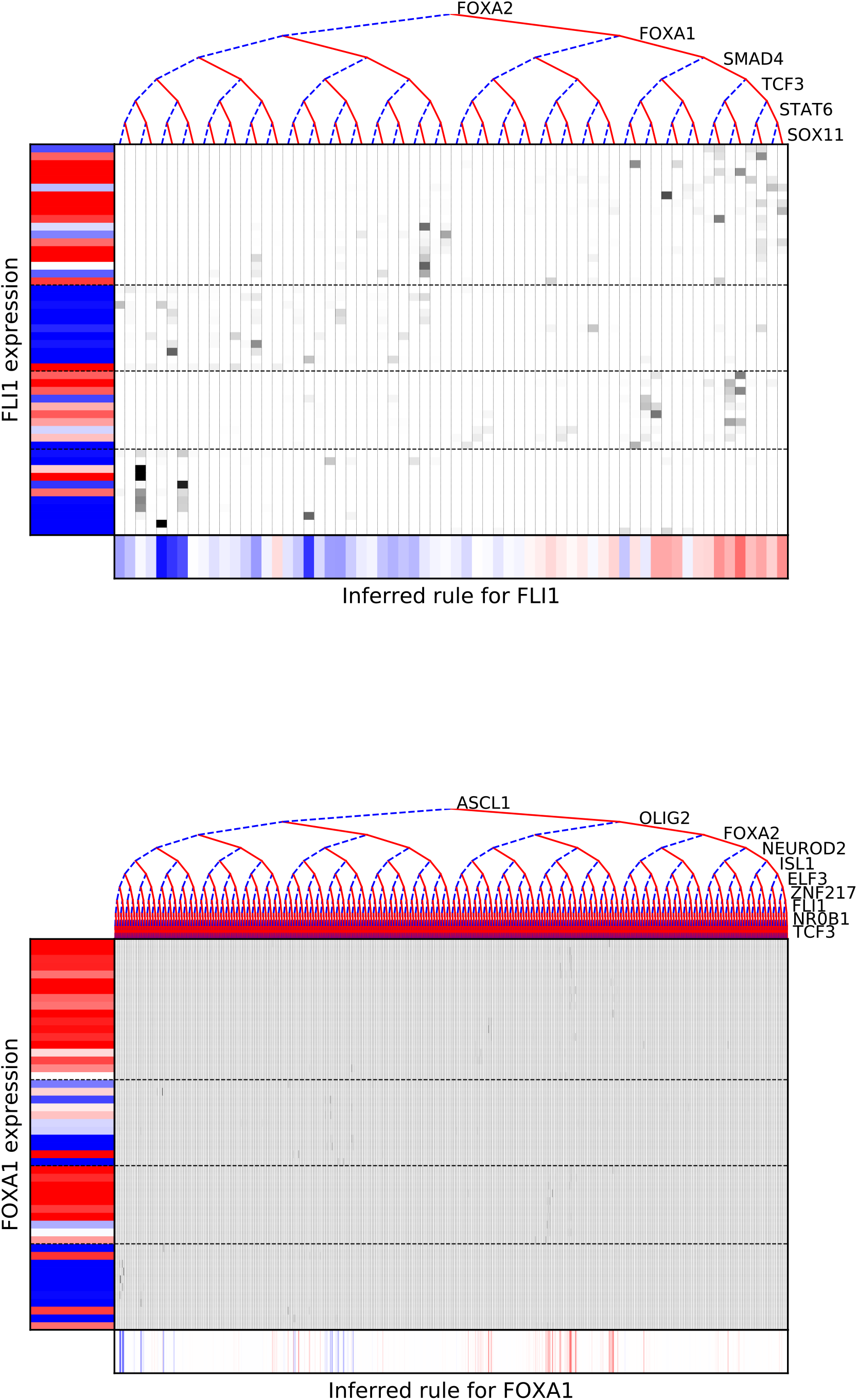

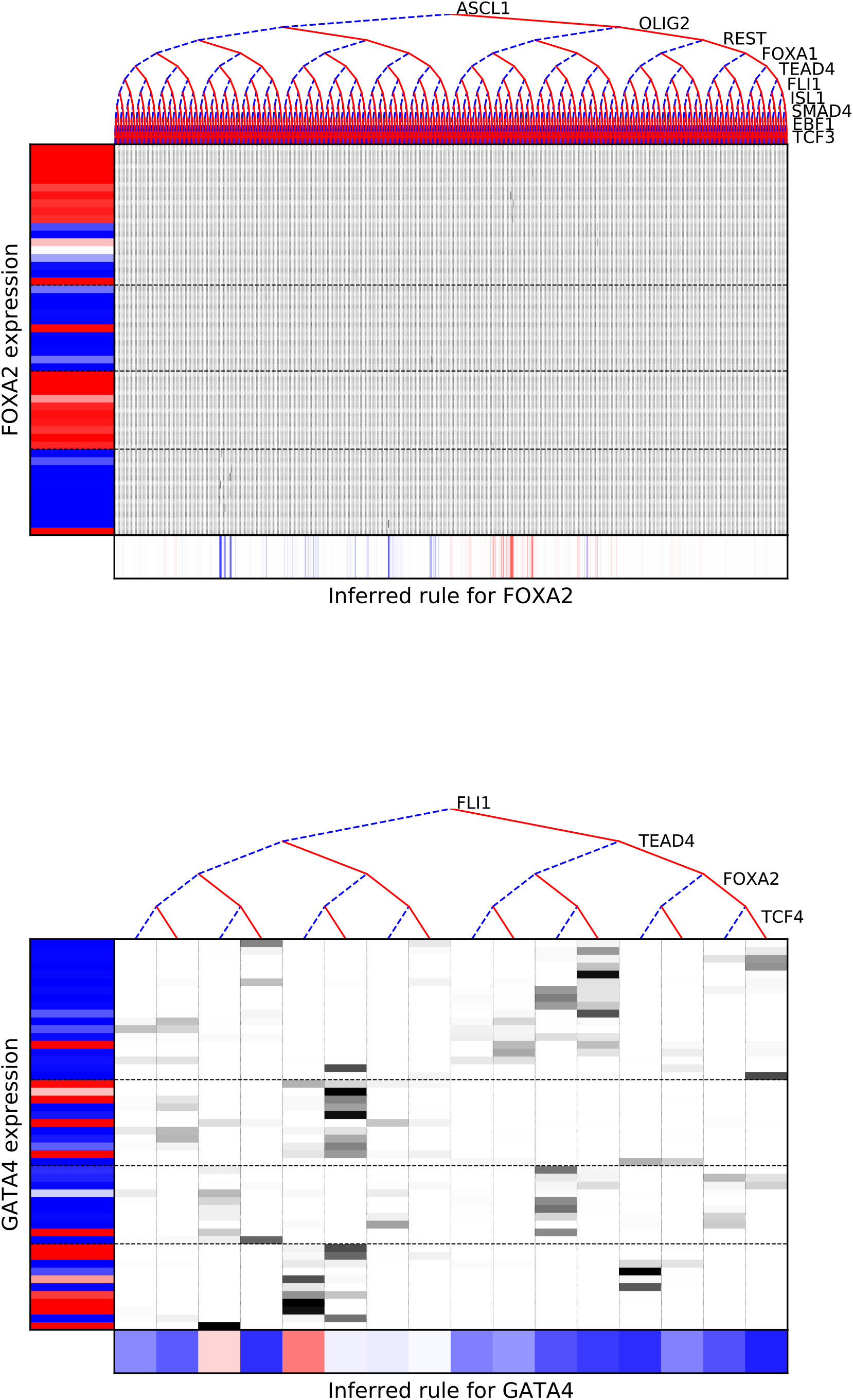

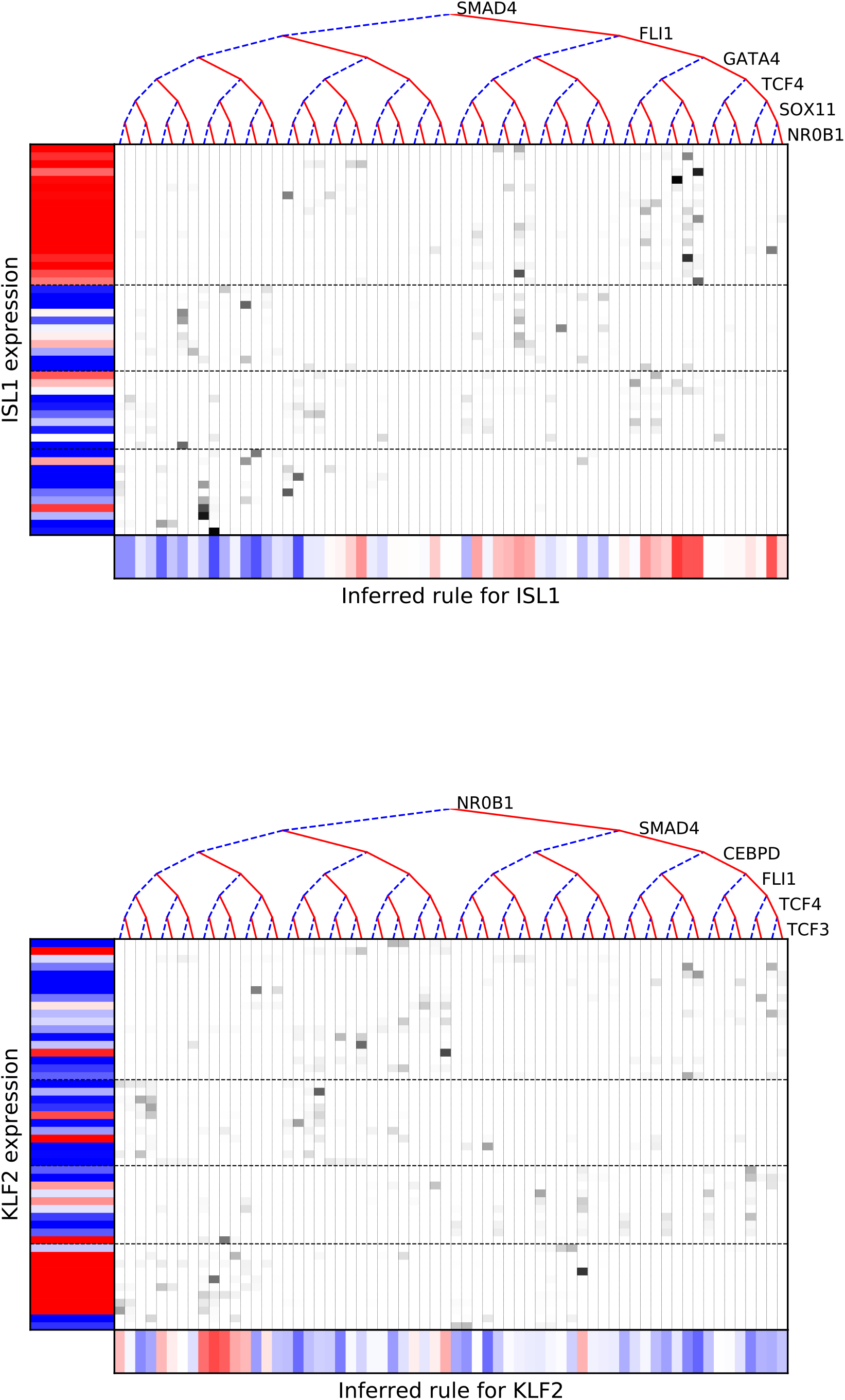

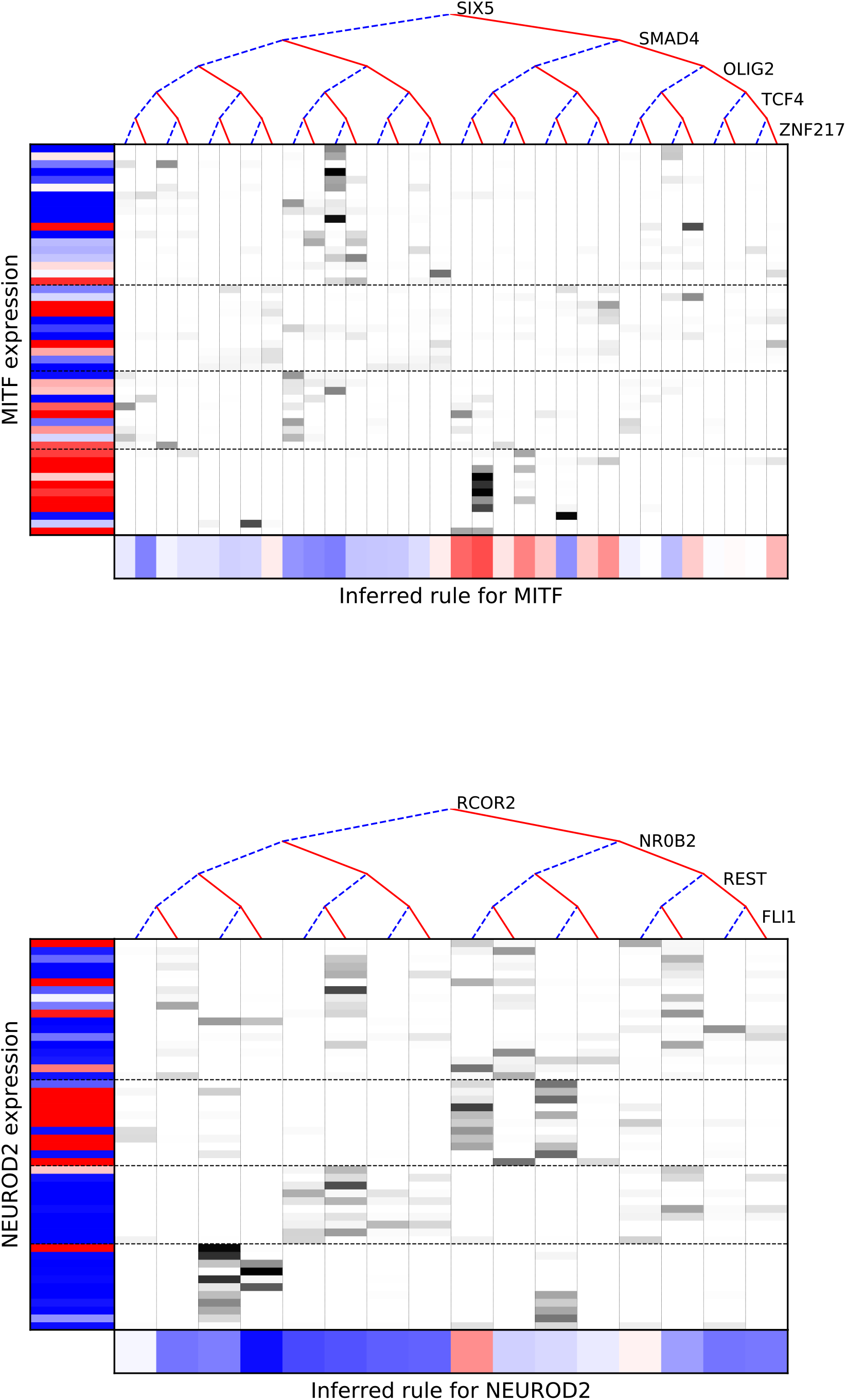

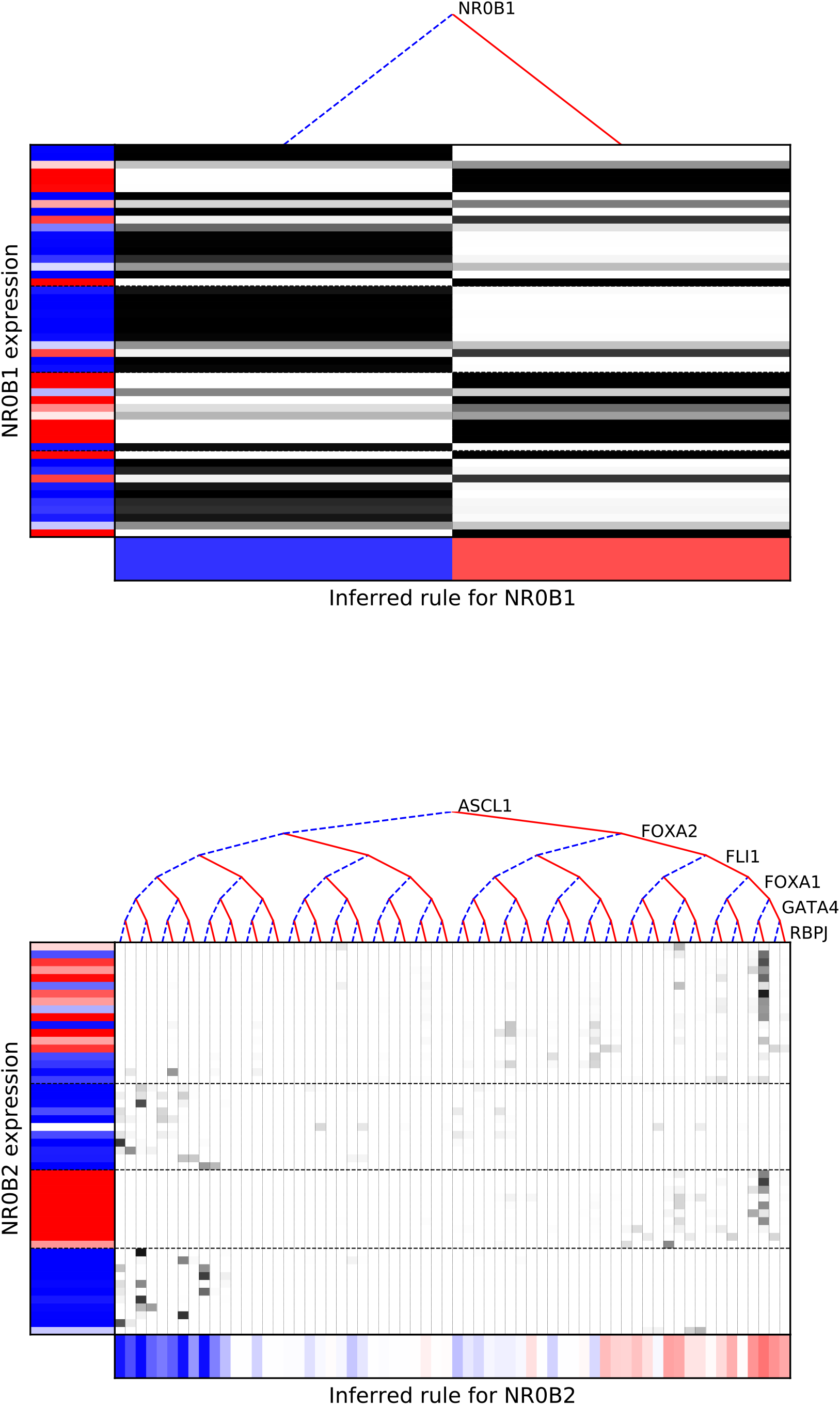

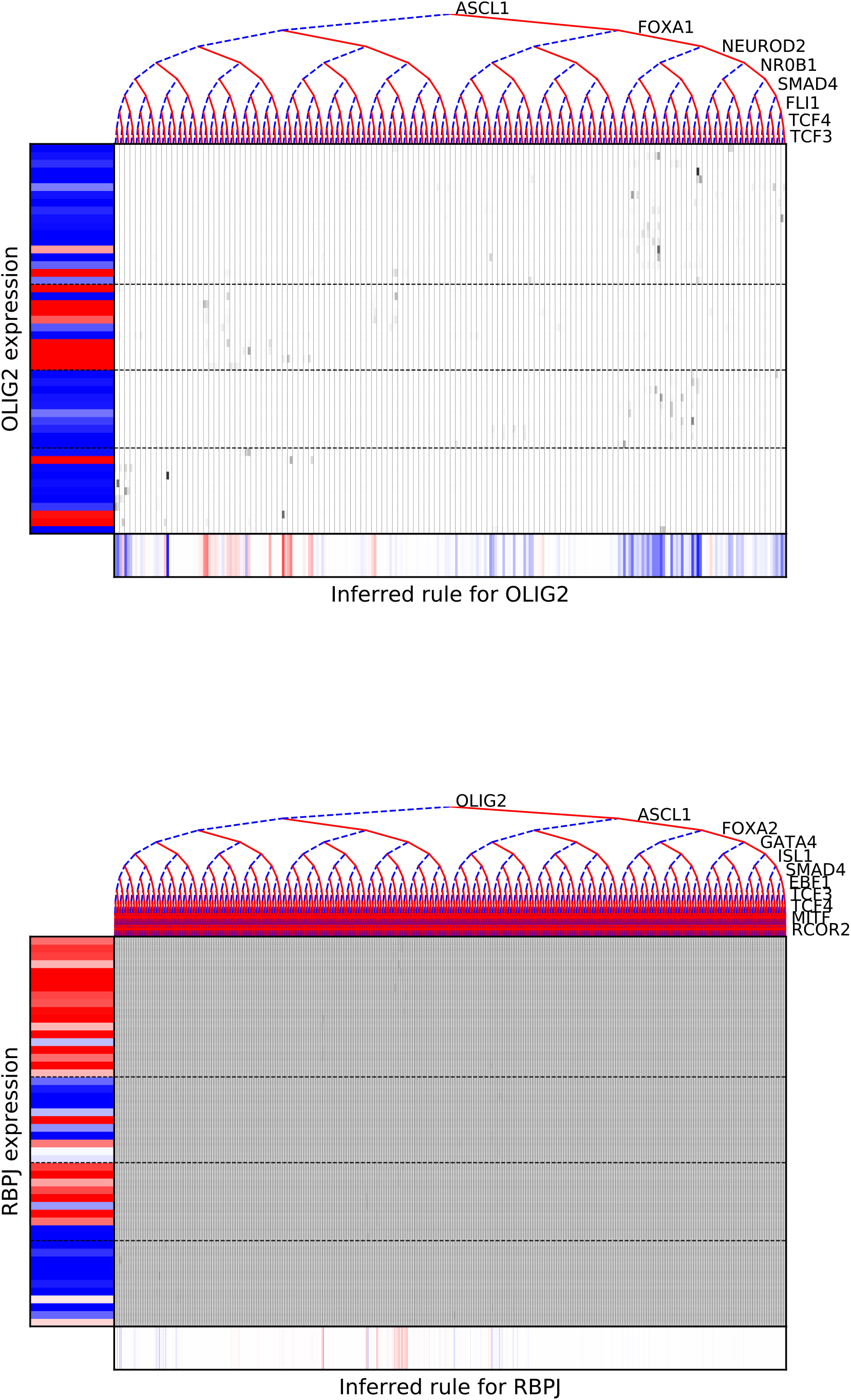

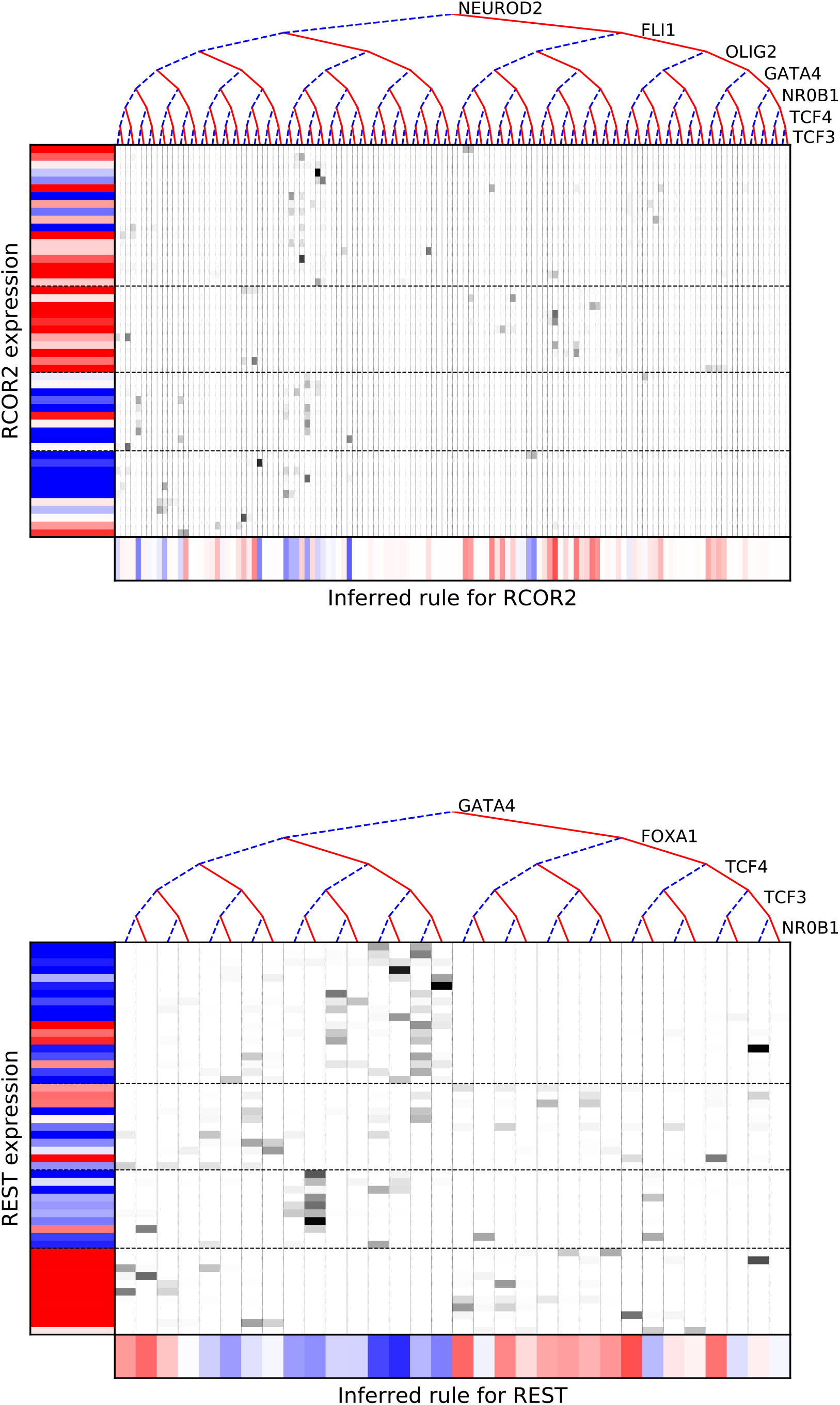

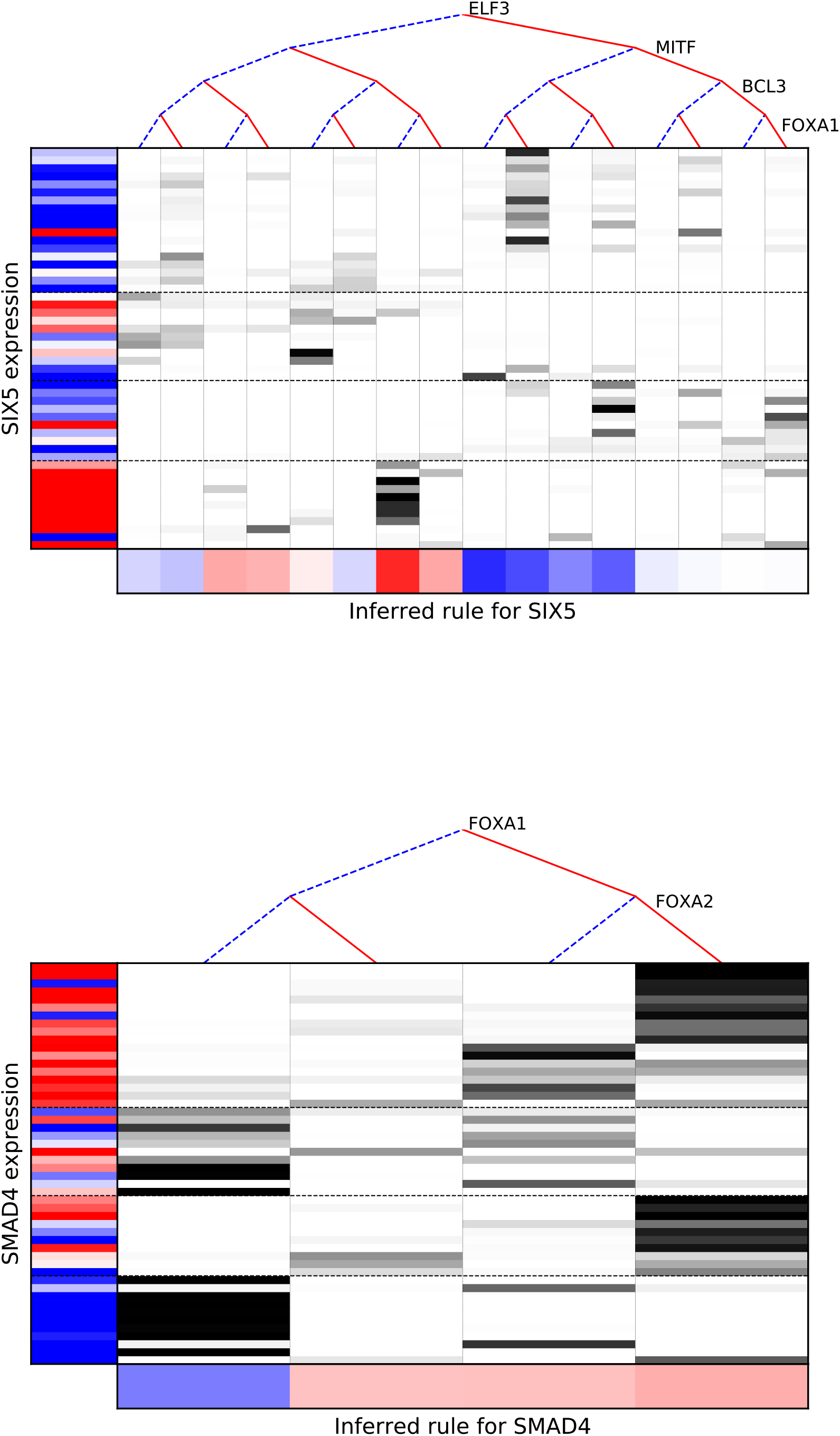

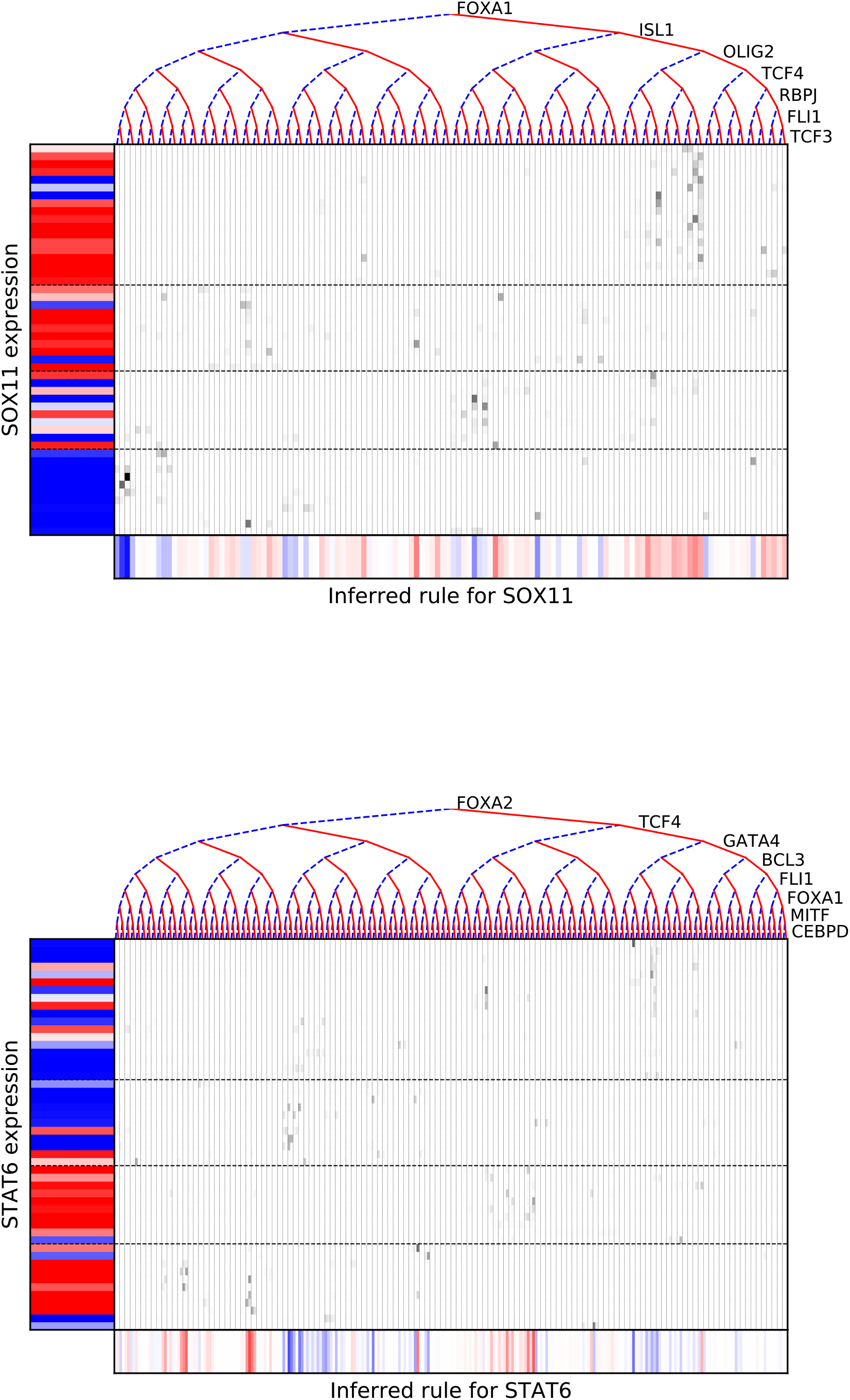

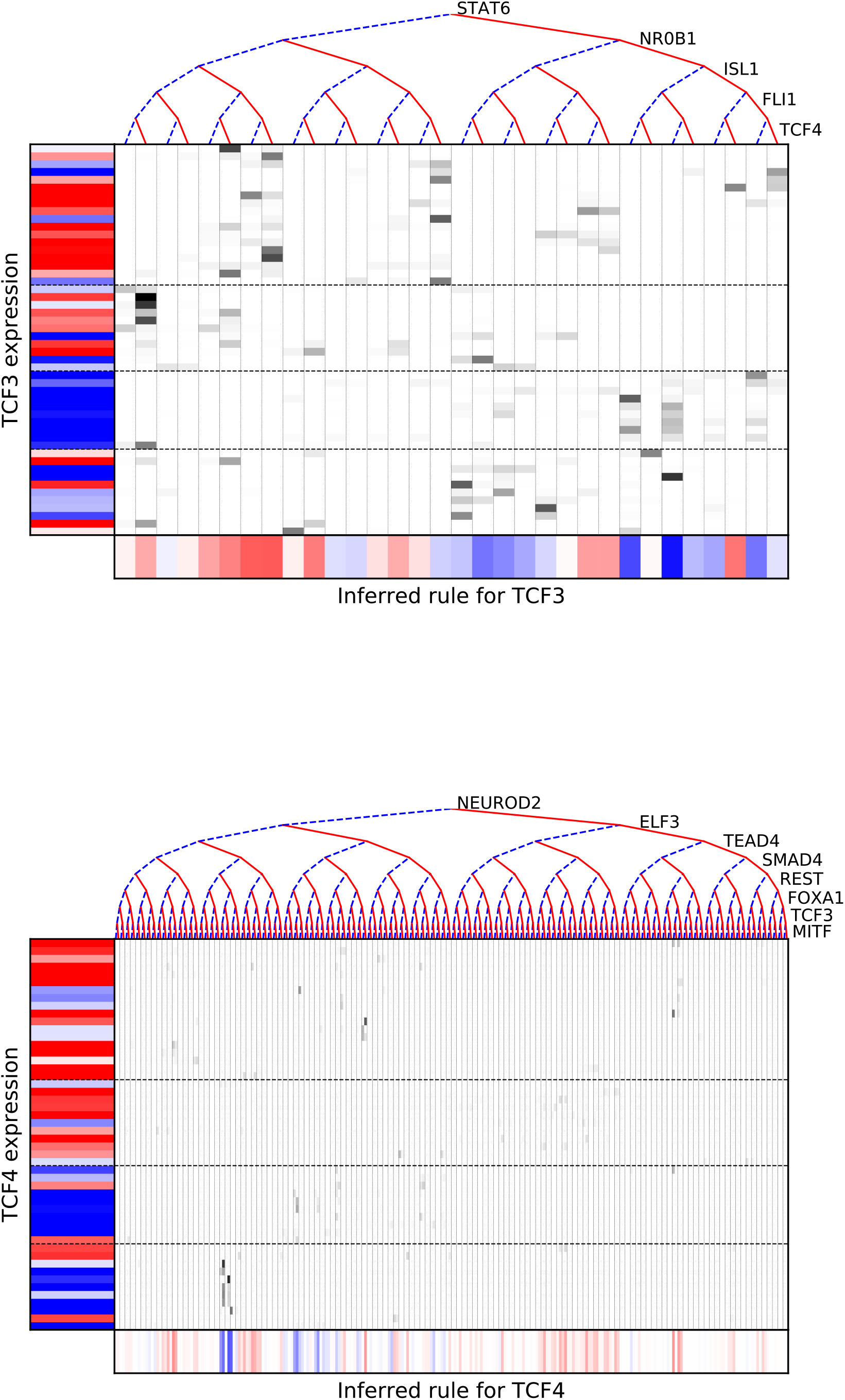

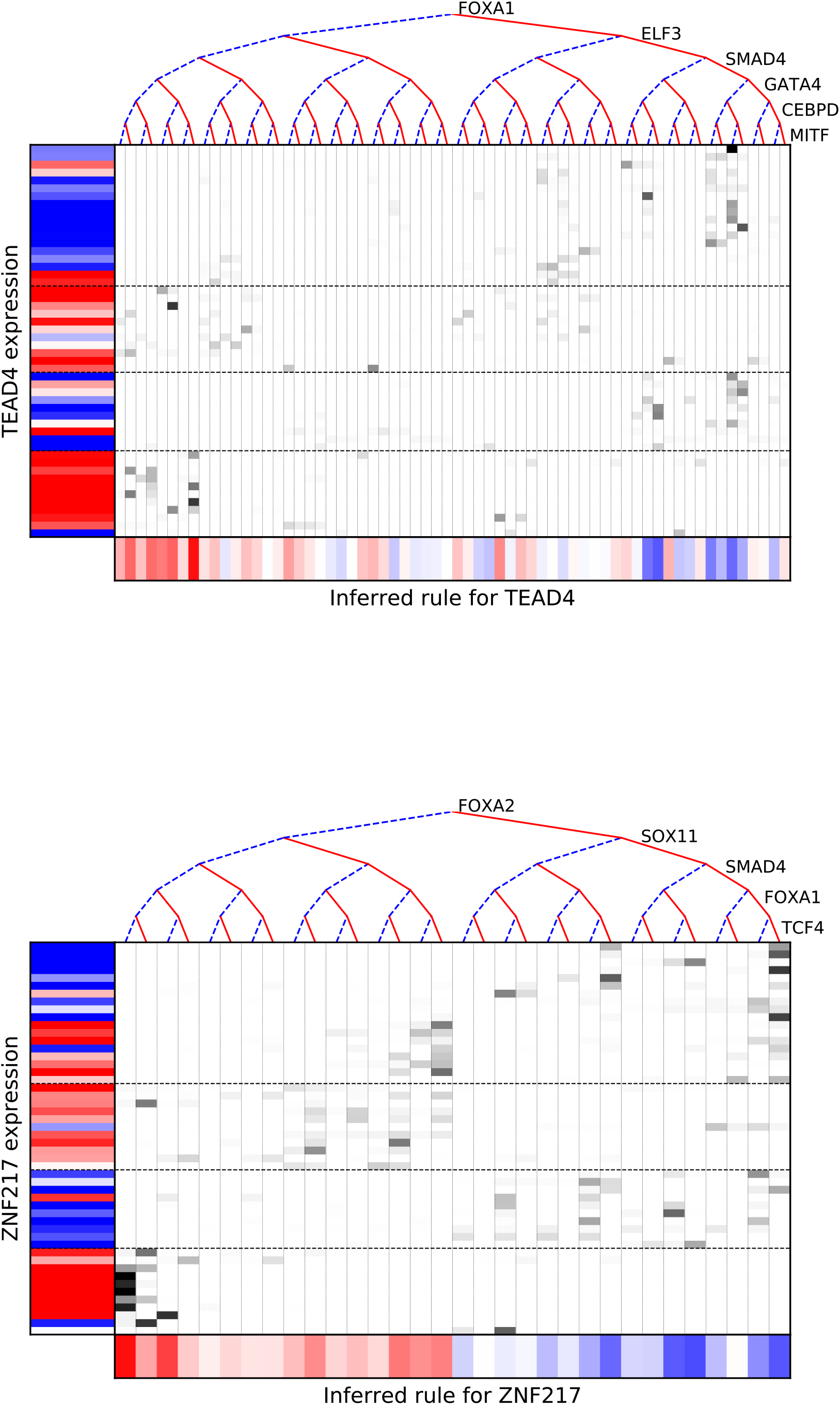

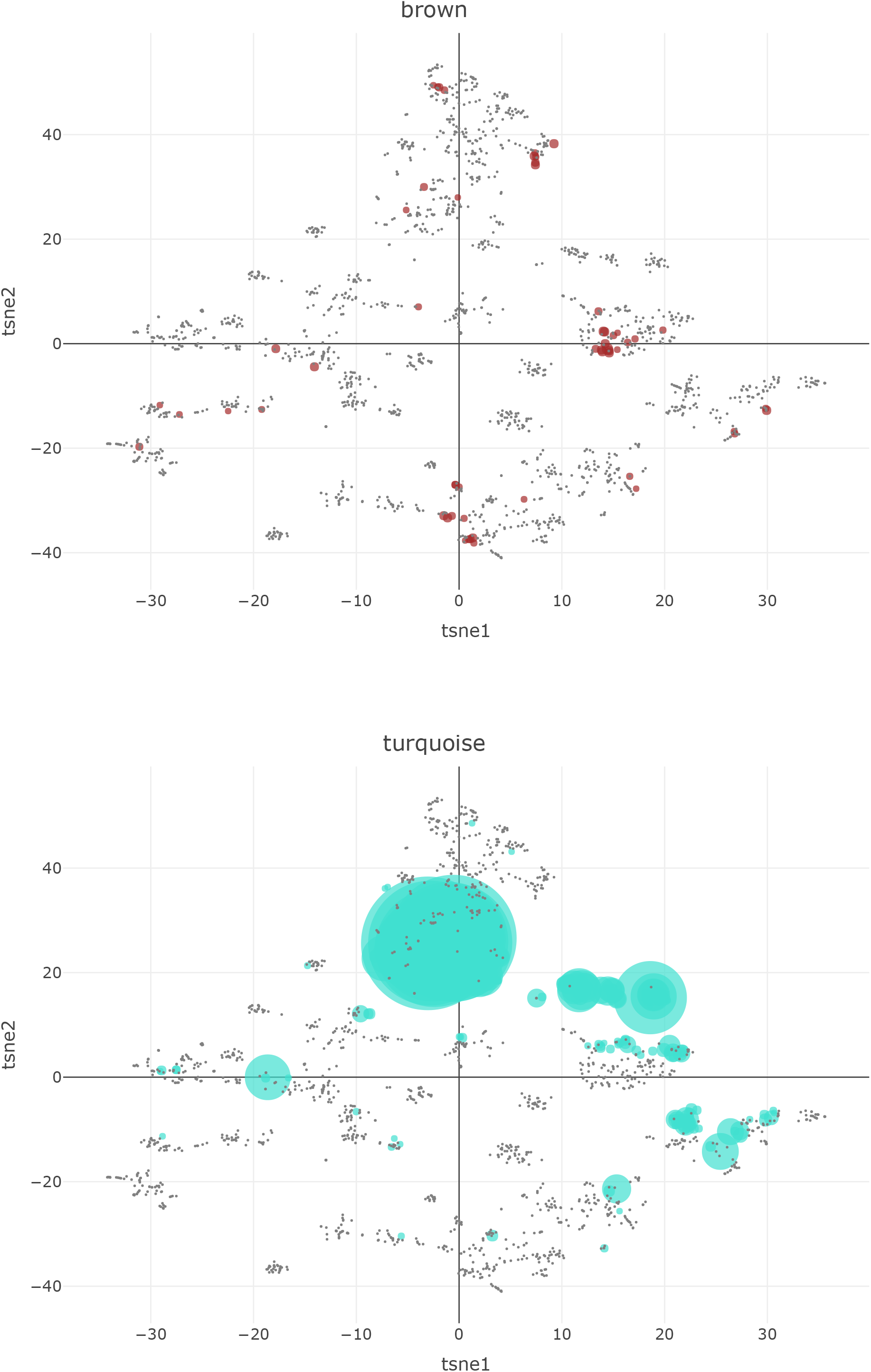

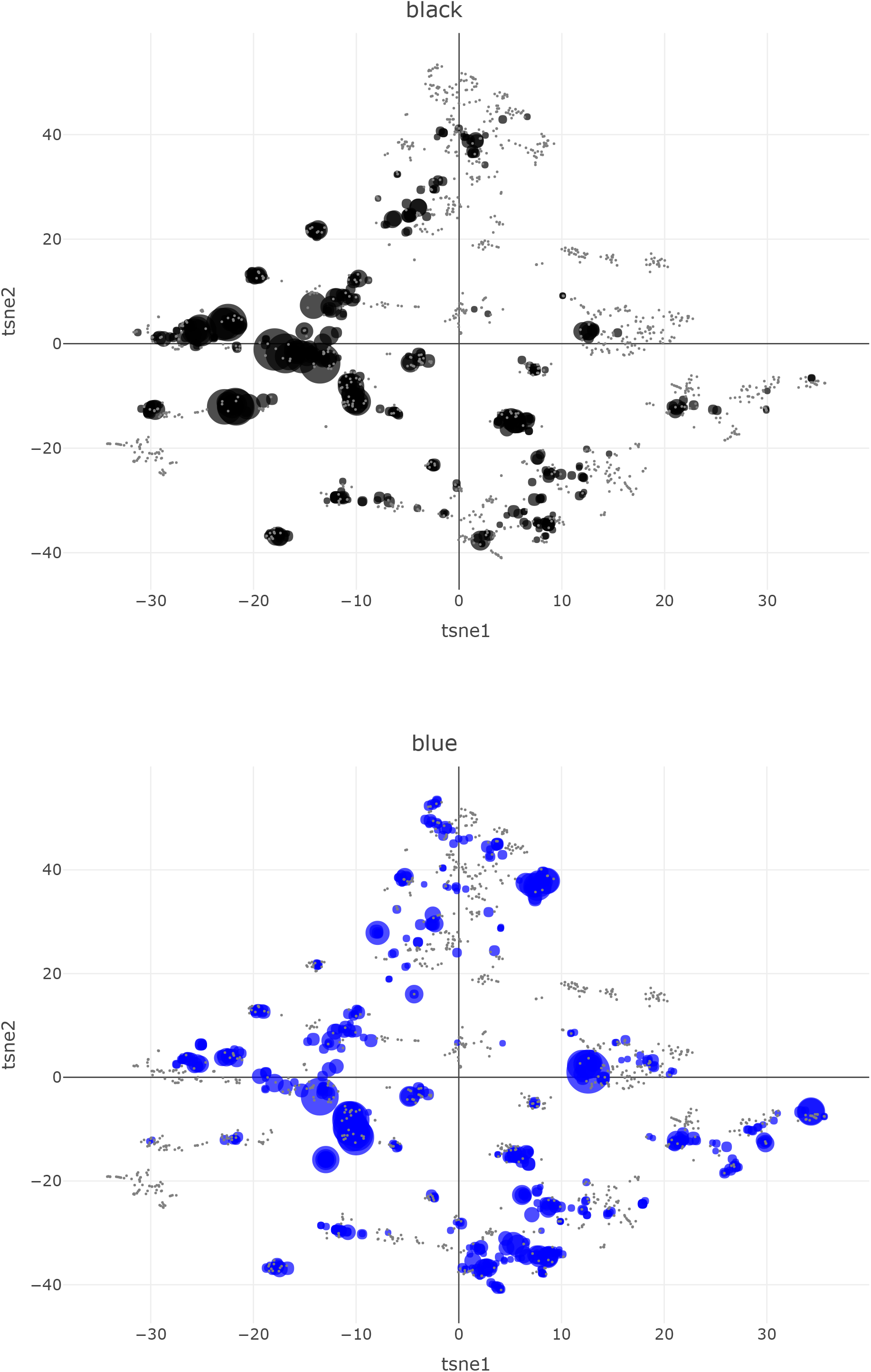

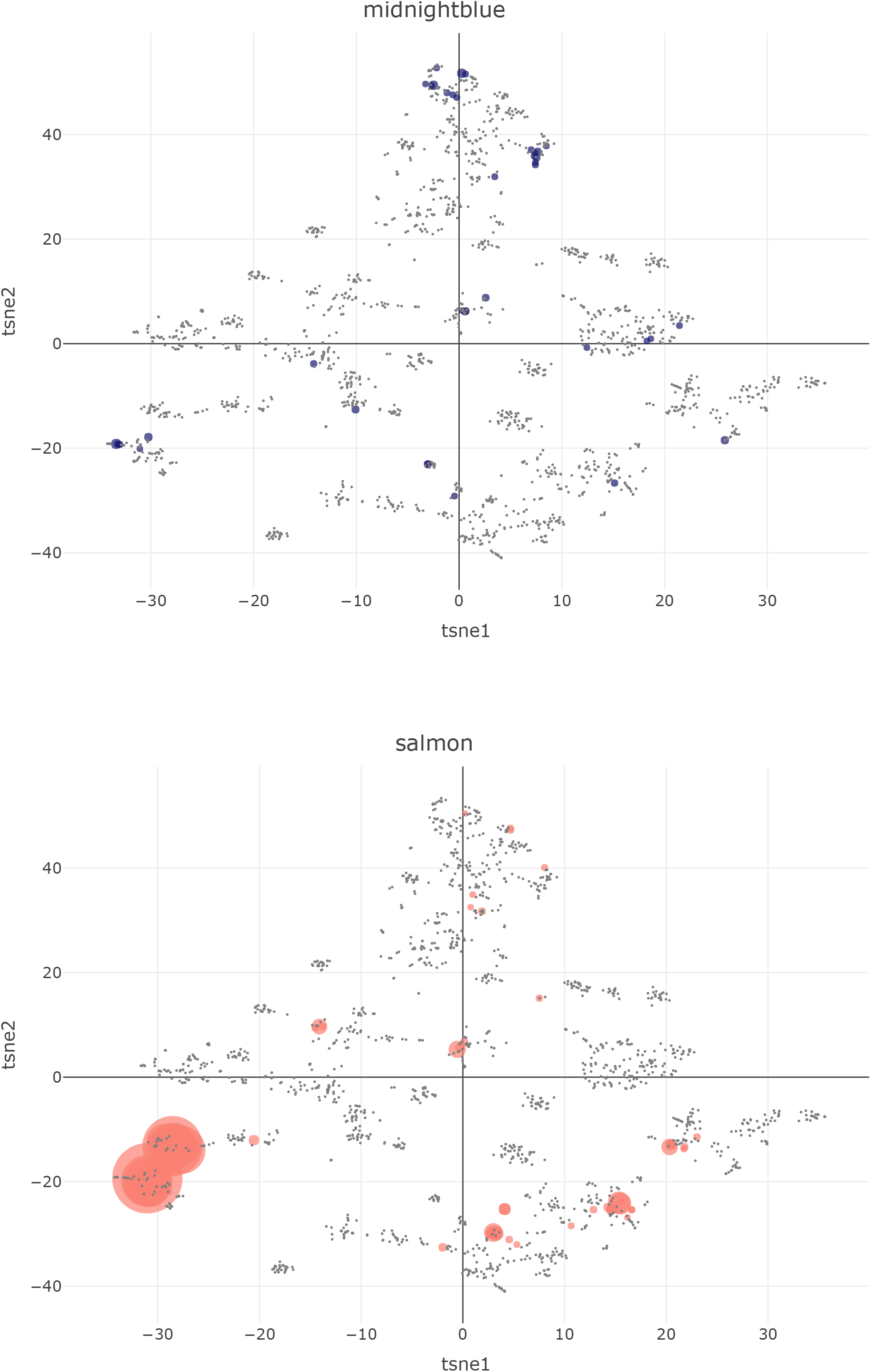

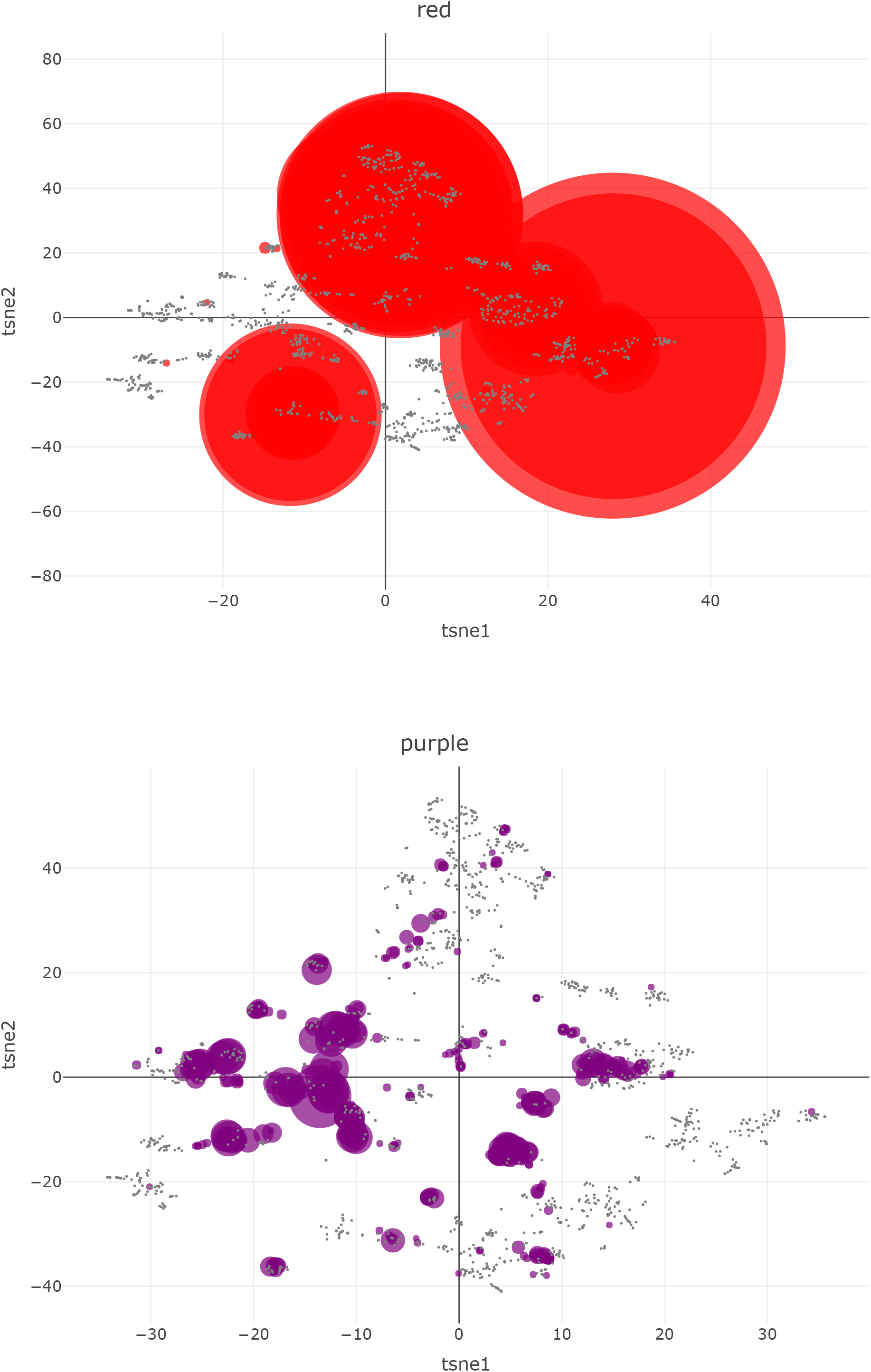

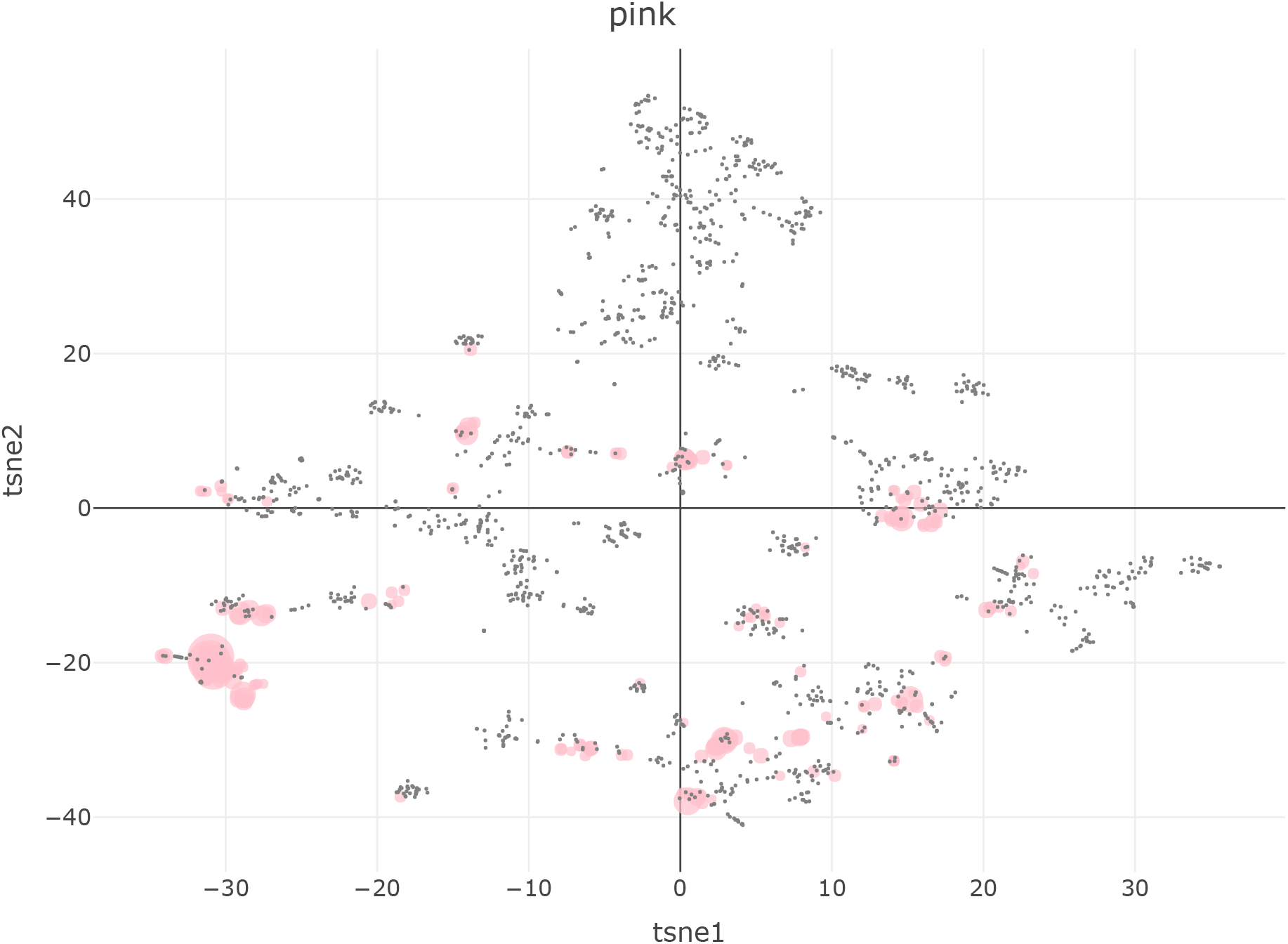

## Acknowledgments

We thank members of the Quaranta laboratory for useful suggestions; members of Carlos Lopez laboratory (Vanderbilt University) for critical feedback and support with computation; and Wade Iams, Christine Lovly, and Jonathan Lehman (Vanderbilt University Medical Center) for their clinical expertise. We also thank the Stanford Functional Genomics Facility for technical support with the 10x Genomics experiments. J.S. is the Harriet and Mary Zelencik Scientist in Children’s Cancer and Blood Diseases.

